# β-III tubulin identifies anti-fibrotic state of pericytes in pulmonary fibrosis

**DOI:** 10.1101/2025.04.15.648984

**Authors:** Ryo Sato, Kosuke Imamura, Tatsuya Tsukui, Teruhiko Yoshida, Yusuke Tomita, Kosuke Fujino, Tokunori Ikeda, Keigo Onizawa, Takumi Sogo, Christian A. Combs, Meera Murgai, Jeffrey B. Kopp, Makoto Suzuki, Takuro Sakagami, Dean Sheppard, Yoh-suke Mukouyama

## Abstract

Pericytes have been implicated in pulmonary fibrosis, yet their activated cellular states and functional roles remain largely unclear. Here, we identified β-III tubulin (Tuj1) as a distinctive marker for fibrosis-associated lung pericytes. Most pericytes in fibrotic regions are Tuj1-positive and interact uniquely with multiple endothelial cells, localizing near collagen-producing fibroblasts and pro-fibrotic SPP1^+^/arginase^+^ macrophages. Tuj1 expression is predominantly induced in pericytes during the fibrotic phase, and Tuj1 gene (*Tubb3*) knockout in mice exacerbates lung fibrosis, accompanied by an increase in the neighboring pro-fibrotic fibroblasts and macrophages, suggesting an anti-fibrotic role for Tuj1-expressing pericytes. Mechanistically, the anti-fibrotic chemokine CXCL10 is upregulated in Tuj1-expressing pericytes, whereas this upregulation is not observed in *Tubb3* knockout. Moreover, CXCL10 inhibits the pro-fibrotic differentiation of macrophages induced by lung fibroblasts in culture, implying that CXCL10 may mediate the anti-fibrotic effects of Tuj1-expressing pericytes. These findings underscore the role of lung pericytes in negatively regulating fibrotic process and their potential as therapeutic targets for pulmonary fibrosis patients.

**Summary:** In response to fibrotic stimuli, lung pericytes upregulate β-III tubulin (Tuj1) expression, adopting an anti-fibrotic phenotype. This phenotype acts as a negative immune regulator of pro-fibrotic macrophage differentiation by releasing CXCL10, thereby counteracting the progression of pulmonary fibrosis.

## Introduction

Pericytes are the mural cells associated with capillaries and small vessels and regulate vascular homeostasis (van Splunder et al., 2024). They maintain vascular integrity, regulate blood flow, and support endothelial cells (ECs). While these traditional roles are well established, emerging evidence has unveiled new pathological aspects of pericyte function that extend beyond their classical vascular roles (van Splunder et al., 2024). Pericytes act as immune hubs to relay inflammatory signals and regulate the extravasation of immune cells (Proebstl et al., 2012; Stark et al., 2013; Duan et al., 2018; Török et al., 2021). In lung diseases, they secrete various cytokines and recruit leukocytes as potential orchestrators of the inflammatory process (Hung et al., 2017; Rayner et al., 2023). These findings highlight a potential therapeutic approach of targeting pericytes and underscore the importance of a comprehensive understanding of these cells in the context of disease progression.

Pericyte activation is prevalent in fibrotic diseases. Pericytes have long been reported to undergo phenotypic alterations upon fibrotic stimuli, acquiring myofibroblast-like characteristics with collagen-producing capacity (Holm et al., 2018; Di Carlo and Peduto, 2018; Barron et al., 2016). However, recent evidence suggests that this transition is tissue-specific (van Splunder et al., 2024). In liver and kidney fibrosis, pericytes significantly contribute to the myofibroblast population (Dobie et al., 2019; Kuppe et al., 2021). In contrast, their involvement in cardiac fibrosis appears to be limited (Kanisicak et al., 2016; Keppe et al., 2022). In pulmonary fibrosis, non-fibroblast cells, including pericytes, contribute minimally to the collagen-producing cell population (Tsukui et al., 2024). These findings highlight the need to define activated pericytes within fibrotic regions and to explore their interactions with pro-fibrotic populations, such as fibroblasts and inflammatory cells.

Recent advances in three-dimensional (3D) imaging combined with pericyte-specific reporter mouse models have demonstrated that lung pericytes exhibit a distinct cellular morphology in vivo, diverging from their conventional morphological characteristics. Lung pericytes extend projections to multiple capillary ECs (Gillich et al., 2021; Klouda et al., 2025), a structural feature also observed in specific organs such as the pancreas (Almaça et al., 2018) and circumventricular organs (Pfau et al., 2024). These 3D imaging techniques and the identification of markers to detect pericyte activation states are necessary for elucidating their roles in pulmonary fibrosis.

Single-cell technologies, especially single-cell RNA sequencing (scRNA-seq), have helped identify activated cellular states and key markers of cells involved in fibrosis, including epithelial cells, fibroblasts, ECs, and macrophages (Kobayashi et al., 2020; Konkimalla et al., 2023; Raslan et al., 2024; Aran et al., 2019). These studies provide novel insights into the cellular landscape of pulmonary fibrosis at single-cell resolution. However, such molecular signatures and cellular states of pericytes during fibrosis progression remain largely unexplored. In this study, leveraging scRNA-seq datasets from the bleomycin-induced pulmonary fibrosis mouse model and 3D imaging visualization, we identified β-III tubulin (Tuj1), which is exclusively expressed in neuronal cells under homeostasis (Janke and Magiera, 2020), as a distinctive marker for activated lung pericyte states in fibrosis. Tuj1 expression is predominantly induced in lung pericytes within fibrotic regions, where Tuj1^+^ pericytes are localized near pro-fibrotic effector cells, including collagen-producing fibroblasts and SPP1^+^/arginase^+^ macrophages. Intriguingly, genetic ablation of Tuj1 gene (*Tubb3*) significantly exacerbated lung fibrosis. Subsequent scRNA-seq analysis of *Tubb3*^-/-^ mice treated with bleomycin demonstrated an expansion of the adjacent pro-fibrotic cell populations, suggesting that Tuj1^+^ pericytes play a negative regulatory role in fibrosis progression. Mechanistically, we found that Tuj1^+^ pericytes exhibit higher expression levels of the anti-fibrotic chemokine CXCL10. Additionally, lung pericytes in *Tubb3*^-/-^ mice displayed significant impairment in CXCL10 upregulation in response to bleomycin-induced injury. In vitro co-culture experiments further revealed that CXCL10 inhibits the pro-fibrotic differentiation of macrophages induced by lung fibroblasts. These findings suggest that in response to fibrotic stimuli, lung pericytes adopt an anti-fibrotic phenotype characterized by Tuj1 expression, acting as a negative immune regulator of pulmonary fibrosis progression.

## Results

### Identification of β-III tubulin (Tuj1) as a marker for activated pericytes in pulmonary fibrosis

To characterize the activated pericyte states in pulmonary fibrosis, we analyzed publicly available scRNA-seq datasets of the bleomycin-induced pulmonary fibrosis mouse model (the bleomycin model). Due to the low abundance of pericytes in the whole lung, we focused on a dataset enriched for mesenchymal cells, which includes a substantial number of pericytes (Tsukui et al., 2020). Throughout this study, we used pericyte-specific markers (e.g., *Higd1b*, *Cox4i2*, *Ndufa4l2*) recently identified through scRNA-seq analysis (Baek et al., 2022) (Fig. S1 A). Re-clustering of the pericyte population yielded 7 distinct clusters (Fig. 1 A). Among these, only cluster 1 consisted exclusively of cells from bleomycin-treated lungs, which we identified as “activated” pericyte populations. This newly emerged activated pericyte subtype exhibited distinct gene expression signatures, including *Rgs5*, a previously identified marker of activated pericyte states (Berger et al., 2005); *Col18a1,* which encodes the angiostatic peptide endostatin; and *Tubb3,* which encodes β-III tubulin (Fig. 1, B and C). Among the upregulated genes, *Tubb3* demonstrated specific expression in lung mural cells, including pericytes and vascular smooth muscle cells (VSMCs), and a minor fraction of fibroblasts, with a more pronounced increase observed in pericytes in response to bleomycin injury (Fig. S1, B and C).

**Figure 1.**
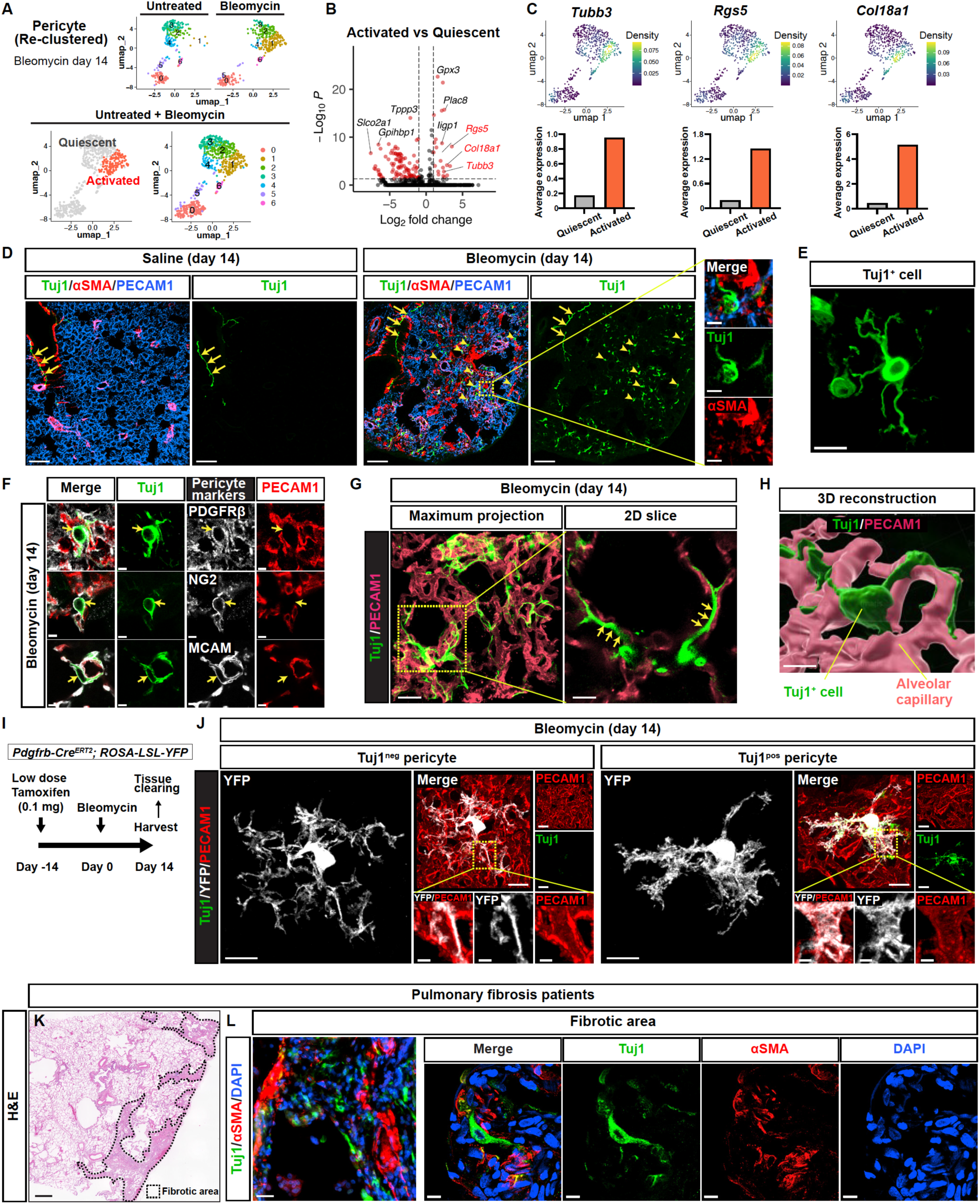
Identification of β-III tubulin (Tuj1) as a marker for activated pericytes in pulmonary fibrosis. **(A−C)** scRNA-seq analysis of the pericyte population from the publicly available dataset of the bleomycin model^21^. **(A)** Uniform manifold approximation and projection (UMAP) plots of the re-clustered pericyte population from untreated and bleomycin-treated lungs. **(B)** Volcano plot showing differentially expressed genes (DEGs) between activated (right) and quiescent (left) pericytes. Red dots represent genes with log2FC > 1 and adjusted *P* < 0.05. **(C)** UMAP plots showing the kernel density estimate for *Tubb3*, *Rgs5*, and *Col18a1* (upper). Bar plots showing the average expression of each gene (lower). **(D)** Immunostaining of saline- or bleomycin-treated lungs with Tuj1 (green), αSMA (red), and PECAM1 (blue). Scale bars, 100 μm; 10 μm (magnified views). **(E)** Tuj1^+^ cells within a fibrotic region at high magnification. Scale bar, 10 μm. **(F)** Immunostaining of bleomycin-treated lungs with pericyte markers (PDGFRβ, NG2, MCAM; gray), Tuj1 (green), and PECAM1 (red). Scale bars, 4 μm. **(G)** Immunostaining of bleomycin-treated lungs with Tuj1 (green) and PECAM1 (pink). Arrows indicate Tuj1^+^ projections aligned with PECAM1^+^ capillary ECs. Scale bars, 20 μm (left); 8 μm (right). **(H)** 3D reconstruction image of Tuj1^+^ cells (green) and PECAM1^+^ alveolar capillary (pink). Scale bar, 7 μm. **(I)** Schematic illustration of mosaic pericyte labeling using *Pdgfrb-Cre^ERT2^; ROSA-LSL-YFP* mice. A single low-dose tamoxifen (0.1 mg) was administered before bleomycin treatment, with lung tissues harvested on day 14 post-treatment. **(J)** Immunostaining of tissue-cleared lungs from bleomycin-treated *Pdgfrb-Cre^ERT2^; ROSA-LSL-YFP* mice with YFP (gray), Tuj1 (green), and PECAM1 (red). Attenuated-maximum intensity projection (attenuated-MIP) was used to visualize 3D lung vascular structures. Boxed regions in attenuated-MIP rendered images of Tuj1^neg^ and Tuj1^pos^ pericytes are magnified as corresponding 3D cropped images on the lower right panels. Scale bars, 10 μm; 2 μm (3D cropped). **(K and L)** Analysis of fibrotic lung tissues resected from lung cancer patients with idiopathic pulmonary fibrosis (IPF). **(K)** Hematoxylin and eosin (H&E) staining of the fibrotic lung tissues. Scale bar, 1000 μm. **(L)** Immunostaining of the fibrotic lung tissues with Tuj1 (green), αSMA (red), and DAPI (blue). Left panel shows a low-magnification image, while right panels present high-magnification images of a representative fibrotic region. Scale bars, 30 μm (left); 10 μm (right).

We next investigated β-III tubulin (Tuj1) protein expression using immunostaining, comparing control lung (saline-treated) with bleomycin-treated lung tissue. Given that Tuj1 is a well-established neuronal marker that labels entire axons (Janke and Magiera, 2020), we observed, as expected, Tuj1 expression in nerves surrounding the αSMA^+^ smooth muscle-covered airways (Fig. 1 D; arrows). Interestingly, bleomycin-treated mice exhibited a marked increase in Tuj1-expressing cells in lung parenchyma (Fig. 1 D; arrowheads, and Fig. 1 E). These Tuj1-expressing cells were negative for αSMA (Fig. 1 D), expressed pericyte markers such as PDGFRβ, NG2, and MCAM (Fig. 1 F), and displayed a highly branched morphology with their projections aligned along PECAM1^+^ capillary endothelial cells (ECs) (Fig. 1, E, G, and H). As previously demonstrated (Gillich et al., 2021; Klouda et al., 2025), our visualization using a mosaic labeling technique with low-dose tamoxifen in *Pdgfrb-Cre^ERT2^; YFP* mice revealed that lung pericytes showed branched cytoplasmic processes interacting with multiple ECs under homeostatic conditions (Fig. S2, A and B). In the bleomycin-treated mice, Tuj1^+^ pericytes displayed thicker projections and exhibited greater coverage of PECAM1^+^ capillary ECs compared to Tuj1^-^ pericytes (Fig. 1, I and J; and Fig. S2 C). Tuj1 expression thus appears to enhance the association of pericytes with ECs.

### Tuj1 is expressed in lung pericytes of human pulmonary fibrosis patients

To assess the clinical relevance of Tuj1 expression, we investigated human fibrotic lung tissues. The previous immunohistochemical study of Tuj1 expression in idiopathic pulmonary fibrosis (IPF) patients has shown that Tuj1 is expressed in pericytes associated with aberrant angiogenesis (Chilosi et al., 2017). Consistent with this observation, our immunohistochemical analysis of the fibrotic lung tissues resected from lung cancer patients with IPF identified Tuj1^+^/αSMA^-^ cells with multiple projections (Fig. 1, K and L), morphologically resembling the Tuj1^+^ pericytes in the bleomycin model. These findings suggest a potential link between Tuj1^+^ pericytes and pulmonary fibrosis in humans.

### Bleomycin injury induces Tuj1 expression predominantly in lung pericytes within fibrotic regions

To examine the spatial distribution of Tuj1^+^ pericytes across the lung, we categorized the bleomycin-treated lung tissue into three distinct regions: airspace, non-fibrotic, and fibrotic areas (Fig. 2, A and B), based on the expression patterns of podoplanin, collagen 1, αSMA, and Hoechst 33342. Podoplanin, a marker for type I alveolar epithelial cells, and Hoechst nuclear staining were used to identify enlarged lung parenchyma. Within these enlarged areas, regions positive for collagen 1 were defined as fibrotic zones. αSMA was used to detect activated fibroblasts, αSMA^+^ smooth muscle-covered airways, and αSMA^+^ VSMC-covered large-diameter blood vessels. These αSMA^+^ airway and vascular structures were excluded from our analysis as they naturally contain collagen under homeostatic conditions, enabling us to focus on bleomycin-induced fibrotic regions. Based on these criteria, we performed machine learning-based area annotation on the bleomycin-treated lung tissue (Fig. 2 B). Density plot analysis revealed that Tuj1^+^ cells localized within the fibrotic regions (Fig. 2 C).

**Figure 2.**
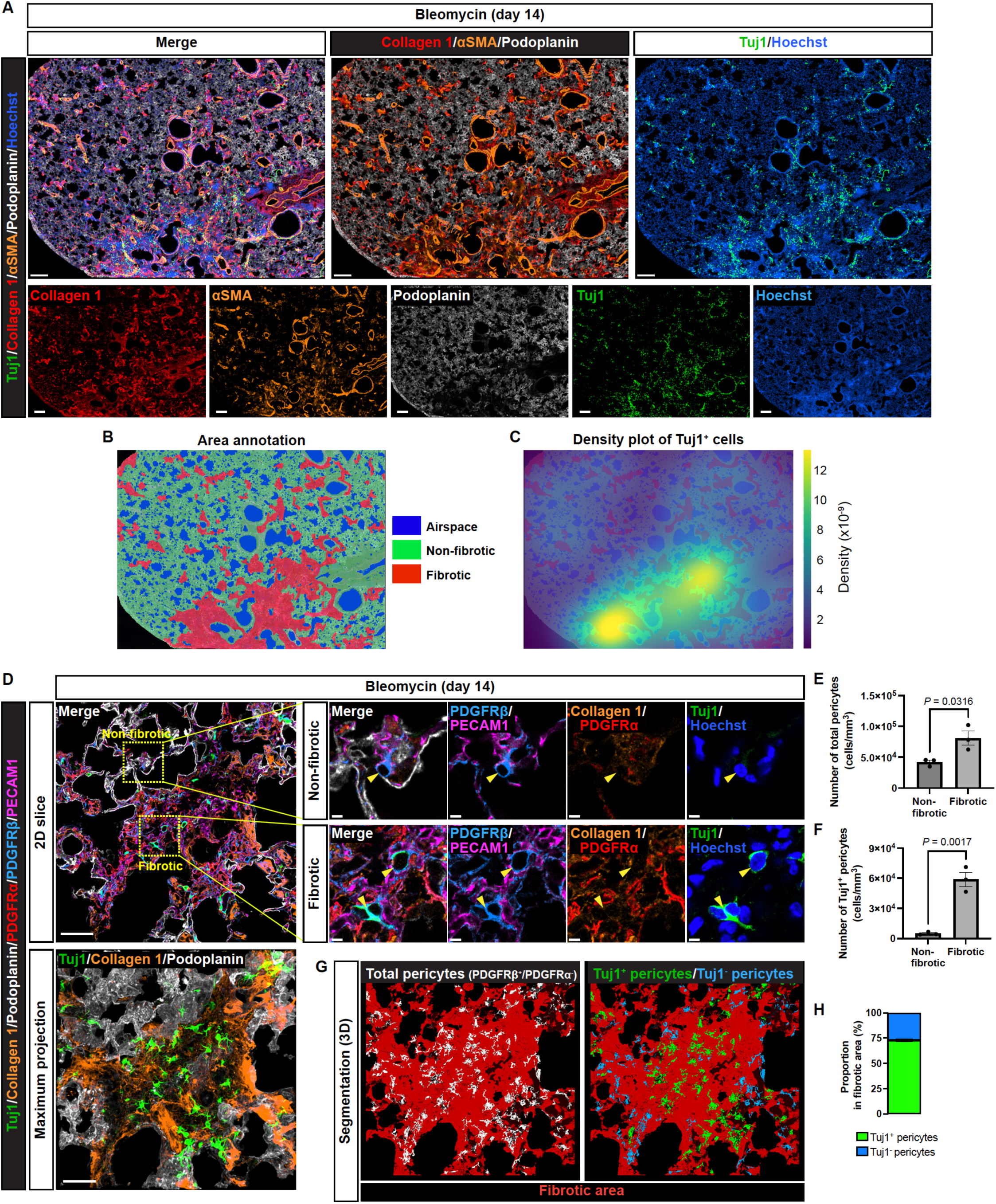
Bleomycin injury induces Tuj1 expression in most lung pericytes exclusively within fibrotic regions. **(A)** Immunostaining of bleomycin-treated lungs with Tuj1 (green), collagen 1 (red), αSMA (brown), podoplanin (gray) and Hoechst33342 (blue). Scale bars, 1000 μm. **(B)** Area annotation image includes airspace (blue), non-fibrotic (green), and fibrotic (red). **(C)** Density plot of Tuj1^+^ cells overlaid on (B). **(D)** Immunostaining of bleomycin-treated lungs labeled with Tuj1 (green), collagen 1 (brown), podoplanin (gray), PDGFRα (red), PDGFRβ (blue), and PECAM1 (magenta). Upper panels show 2D slice images. Boxed regions in non-fibrotic and fibrotic areas in the left panel are magnified in the right panels. Arrowheads indicate pericytes. Lower left panel shows a maximum intensity projection image. Scale bars, 50 μm (upper and lower left panels); 5 μm (upper right panels). **(E and F)** Quantification of the total number of pericytes (E) and Tuj1^+^ pericytes (F) within non-fibrotic and fibrotic areas (*n* = 3 mice, mean ± SEM, unpaired *t* test). **(G)** 3D segmentation of total lung pericytes (gray), Tuj1^+^ pericytes (green) and Tuj1^-^ pericytes (blue) within fibrotic areas (brown) from the immunofluorescence image in (C). **(H)** Proportion of Tuj1^+^ and Tuj1^-^ pericytes within the fibrotic area (mean ± SEM).

We then examined whether the majority of lung pericytes in the fibrotic region expressed Tuj1. To address this, we employed a combinatorial approach to identify lung pericytes, defining them as PDGFRβ^+^/PDGFRα^-^ cells closely associated with PECAM1^+^ capillary ECs. This approach allowed us to clearly visualize lung pericytes and distinguish between Tuj1^+^ and Tuj1^-^ subpopulations (Fig. 2 D). In non-fibrotic areas, lung pericytes showed no Tuj1 expression, whereas pericytes in fibrotic regions exhibited high Tuj1 expression (Fig. 2 D) and were significantly increased in number (Fig. 2 E). Tuj1-positive pericytes were exclusively localized to fibrotic regions (Fig. 2, D and F), where approximately 75% of pericytes expressed Tuj1 (Fig. 2, G and H).

Since we observed abundant pericytes within fibrotic regions compared to non-fibrotic regions (Fig. 2 E), we next examined the proliferation state of lung pericytes in both regions (Fig. 3 A). Our quantification analysis showed an increased expression of the proliferation marker Ki67 in lung pericytes within the fibrotic regions (Fig. 3 B). Interestingly, when focusing on the fibrotic regions, we found no significant difference in Ki67 expression between Tuj1^+^ and Tuj1^-^ pericytes (Fig. 3 C), suggesting that Tuj1 expression does not directly influence pericyte proliferation in this context. Since pericyte proliferation is closely coordinated with ECs during pathological angiogenesis, we examined the expression of TrkB (also known as Ntrk2, neurotrophic receptor tyrosine kinase 2) in PECAM1^+^ capillary ECs (Fig. 3 D), which has recently been identified as a marker of the activated EC signature linked to angiogenesis (Raslan et al., 2024). Indeed, we observed a close association between Tuj1^+^ pericytes and TrkB^+^ capillary ECs (Fig. 3 D), suggesting that Tuj1^+^ pericytes predominantly localize to fibrotic regions undergoing angiogenesis.

**Figure 3.**
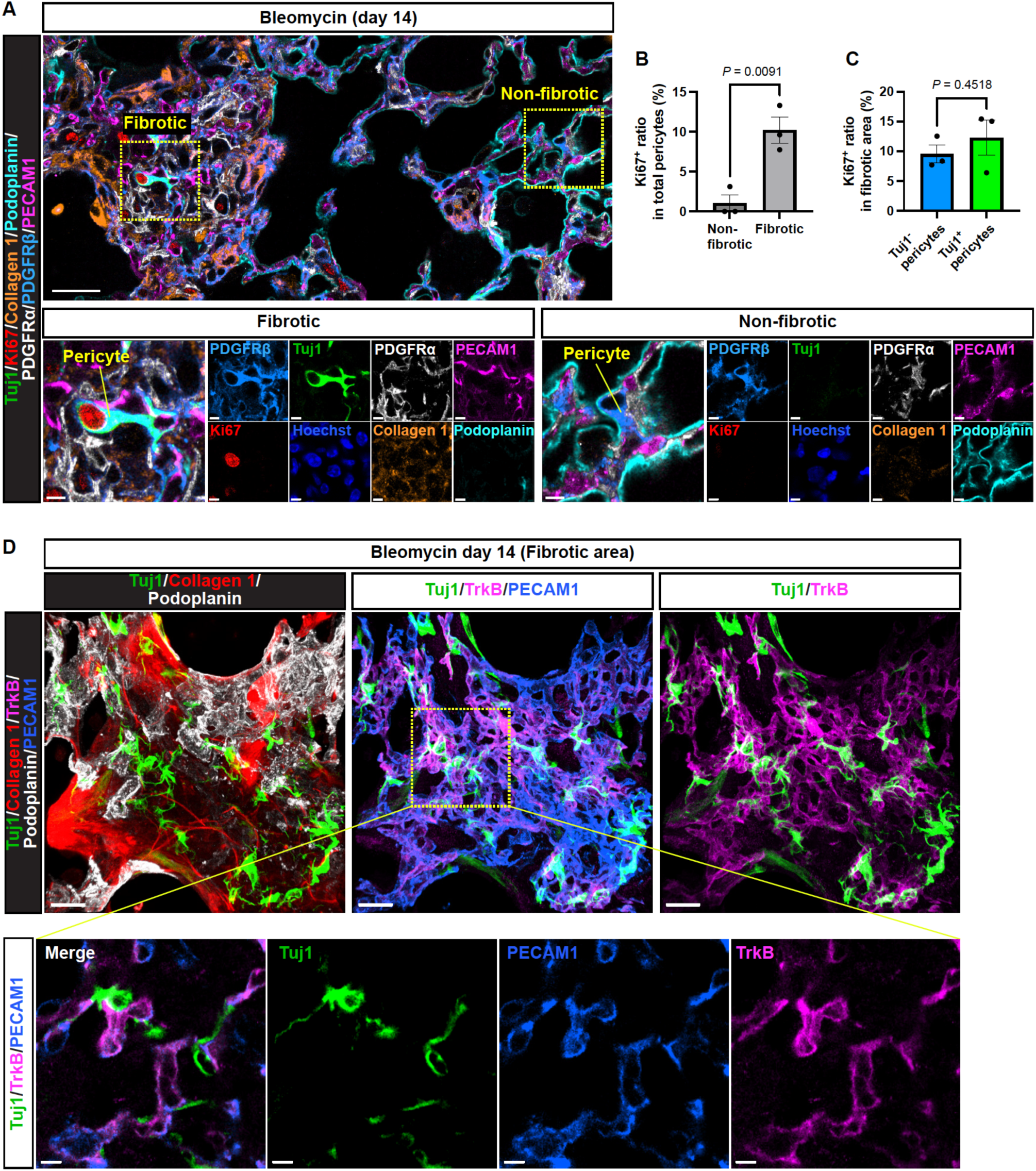
Tuj1^+^ pericytes associate with TrkB^+^ activated capillary endothelial cells. **(A)** Immunostaining of bleomycin-treated lungs (single z-slice) with Tuj1 (green), Ki67 (red), collagen 1 (brown), podoplanin (cyan), PDGFRα (gray), PDGFRβ (blue), and PECAM1 (magenta). Scale bars, 30 μm (upper); 5 μm (lower). **(B)** Quantification of the Ki67-positive ratio in total pericytes within non-fibrotic and fibrotic areas (*n* = 3 mice, mean ± SEM, unpaired *t* test). **(C)** Quantification of Ki67-positive pericytes as a percentage of Tuj1^-^ and Tuj1^+^ pericyte populations in fibrotic areas (*n* = 3 mice, mean ± SEM, unpaired *t* test). **(D)** Immunostaining of bleomycin-treated lungs with Tuj1 (green), collagen 1 (red), TrkB (magenta), podoplanin (gray), and PECAM1 (blue). Upper: maximum projection images; Lower: magnified single slice images. Scale bars, 20 μm (upper); 5 μm (lower).

Although the scRNA-seq analysis detected *Tubb3* RNA expression in lung mural cells (pericytes and VSMCs) and a minor fraction of fibroblasts (Fig. S1 B), our immunostaining analysis revealed that Tuj1 protein expression was predominantly observed in αSMA-negative pericytes in bleomycin-treated lungs. We employed *Col1a1-EGFP* (*Col-GFP*) reporter mice, which enabled clear visualization of collagen-expressing fibroblasts (Fig. 4 A). In the fibrotic lung regions, Col-GFP^high^/αSMA^+^ fibroblasts were abundant; however, their Tuj1 expression levels were negligible or markedly low (Fig. 4, A and B; Fibroblast). In contrast, Tuj1^+^ cells in these regions were predominantly Col-GFP^low^ ^or^ ^negative^/αSMA^-^ pericytes (approximately 80%) rather than Col-GFP^low or negative^/αSMA^high^ VSMCs (around 20%) (Fig. 4, A−C; Pericyte and VSMC). Taken together, these data indicate that Tuj1 expression is predominantly induced in pericytes within fibrotic lung regions in response to bleomycin injury.

**Figure 4.**
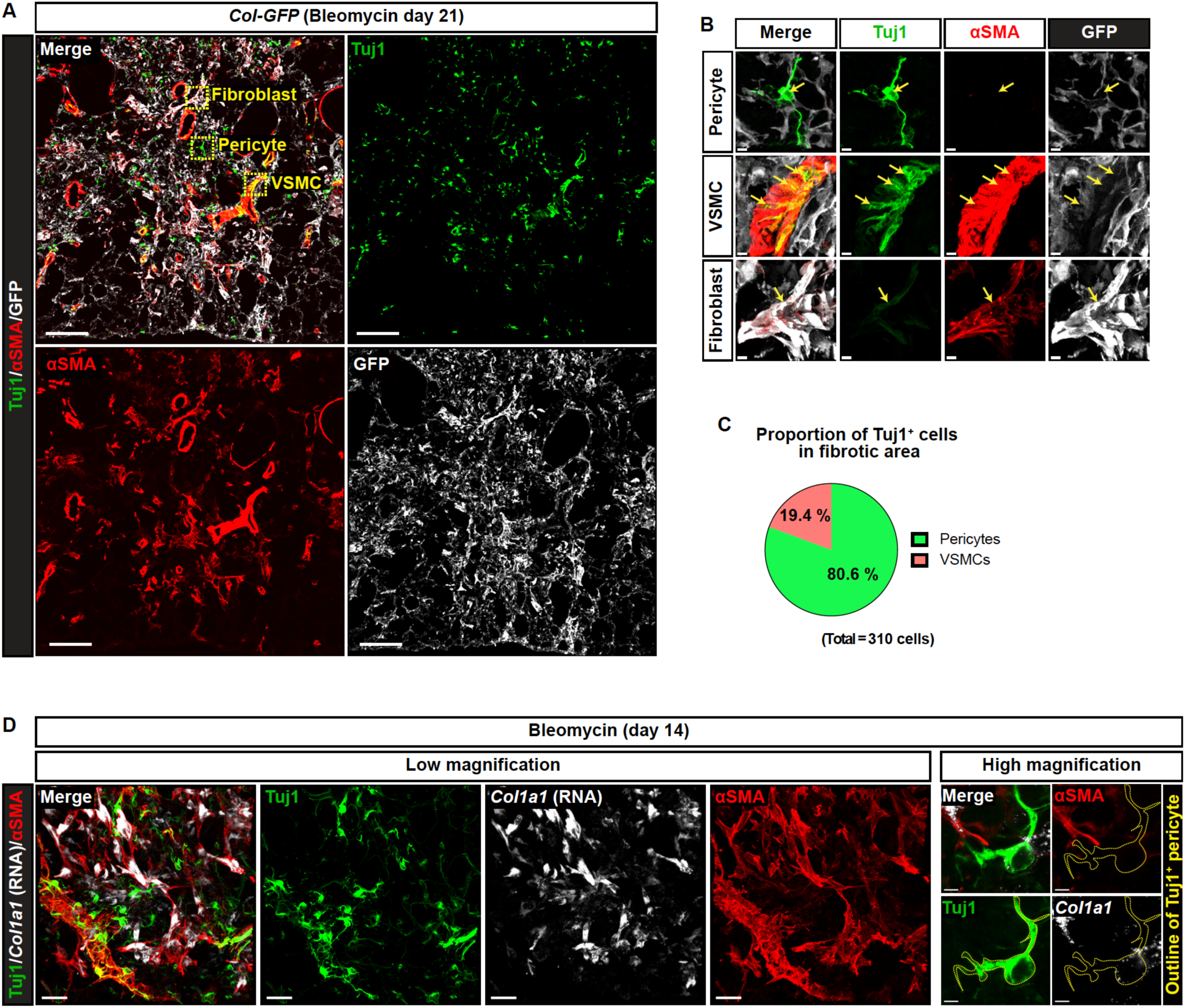
Tuj1^+^ pericytes are not collagen-producing cells. **(A)** Immunostaining of bleomycin-induced fibrotic lungs (day 21) from *Col1a1-EGFP* (Col-GFP) reporter mice with Tuj1 (green), αSMA (red), and GFP (gray). Boxed regions are magnified in the panels in (B) to highlight Tuj1-expressing pericyte, VSMC, and fibroblast. Scale bar, 150 μm. **(B)** Magnified images of Tuj1^+^ pericytes (αSMA^-^, GFP^-^; upper panels), Tuj1^+^ VSMCs (αSMA^+^, GFP^-^; middle panels), and Tuj1^weak^ fibroblasts (αSMA^+^, GFP^+^; lower panels). Scale bars, 5 μm. **(C)** Proportion of Tuj1^+^ pericytes and VSMCs within total Tuj1^+^ cells in fibrotic area. **(D)** RNA in situ hybridization combined with immunostaining of bleomycin-treated lungs, labeled with *Col1a1* mRNA (gray), Tuj1 (green), and αSMA (red). Left: lower-magnification overview images of fibrotic area. Right: higher-magnification images of Tuj1^+^ pericytes. Scale bars, 30 μm (left); 3 μm (right).

### Tuj1^+^ pericytes are not collagen-producing cells

While lung pericytes have been reported to acquire collagen-producing myofibroblast-like characteristics during fibrosis (Di Carlo and Peduto, 2018; Barron et al., 2016), the analysis using *Col-GFP* reporter mice revealed that Tuj1^+^ pericytes showed low levels of Col-GFP expression (Fig. 4, A and B). Supporting this finding, RNA *in situ* hybridization for *Col1a1* gene, combined with Tuj1 and αSMA immunostaining, showed that αSMA^+^ fibroblasts express *Col1a1,* while Tuj1^+^ pericytes do not (Fig. 4 D). These results suggest that Tuj1^+^ pericytes are not collagen-producing cells in the fibrotic regions, highlighting a potentially distinct role for these cells in the fibrotic process.

### Tissue-resident lung pericytes upregulate Tuj1 expression in response to fibrotic stimuli

We investigated the origin of Tuj1^+^ pericytes by conducting a series of lineage-tracing experiments. Utilizing the *Pdgfrb-Cre^ERT2^*driver with *Cre*-dependent *ROSA-LSL-YFP* reporter to label all mural cells, we observed that Tuj1^+^ pericytes originated from this lineage, as demonstrated by the co-expression of the YFP reporter (Fig. 5, A−D; 90.3 ± 0.88%). In contrast, the *αSMA-Cre^ERT2^*driver with *Cre*-dependent *ROSA-LSL-RFP* reporter, which specifically labels smooth muscle cells, did not mark Tuj1^+^ branched pericytes (Fig. 5, A−C; 0%). This finding aligns with our immunostaining results, showing that Tuj1^+^ branched pericytes are negative for αSMA. However, a small number of Tuj1^+^/αSMA^+^ VSMCs were observed in large-diameter blood vessels covered by VSMCs and were labeled by both the *Pdgfrb-Cre^ERT2^* and *aSMA-Cre^ERT2^*drivers (Fig. S3, A and B; 86.7 ± 2.33% and 80 ± 10.07%, respectively). This distinct labeling pattern between pericytes and VSMCs indicates that these cell types maintain separate identities without undergoing transdifferentiation. Additional lineage-tracing experiments using the *NG2-Cre^ERT2^* driver with *Cre*-dependent *ROSA-LSL-YFP* reporter supported the finding that Tuj1^+^ pericytes derive from lung pericytes (Fig. S3 C). Notably, as shown in the aforementioned studies (Fig. 2, G and H), most *Pdgfrb*-lineage pericytes expressed Tuj1 in the collagen 1^high^ fibrotic regions (Fig. 5 D), with Tuj1-expressing states representing the majority of pericytes in these areas. To exclude a fibroblastic source of Tuj1^+^ pericytes, we used the *Scube2-Cre^ERT2^* and *Cthrc1-Cre^ERT2^* drivers with *Cre*-dependent *ROSA-LSL-tdTomato* reporter to label alveolar and fibrotic lung fibroblasts, respectively (Tsukui et al., 2024). We found no contribution from either fibroblast population to the Tuj1^+^ pericytes (Fig. S3, D and E). Taken together, these findings indicate that tissue-resident pericytes in the lung upregulate Tuj1 expression in response to fibrotic stimuli, but the progeny of normal alveolar fibroblasts and fibrotic fibroblasts do not.

**Figure 5.**
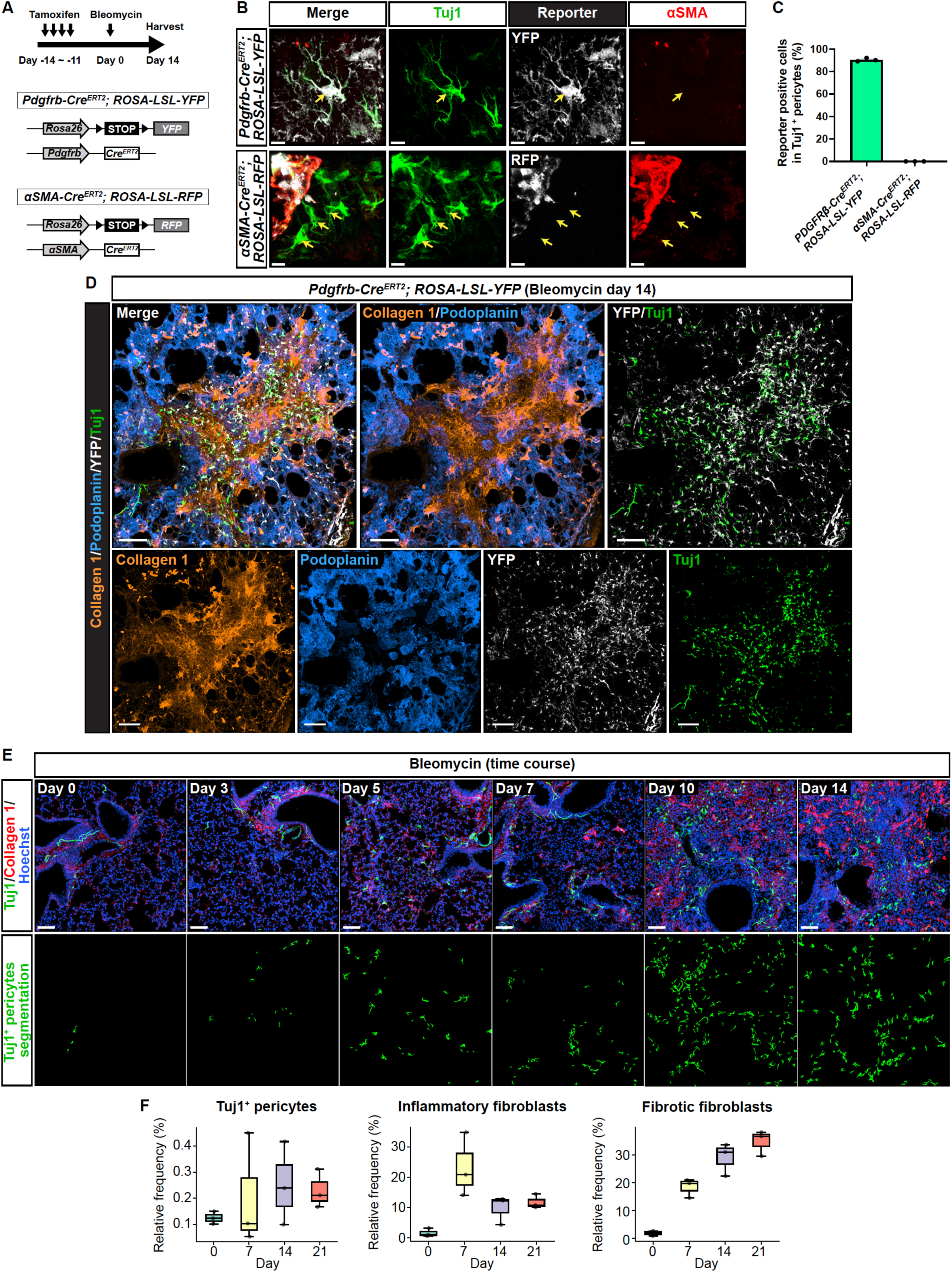
Tuj1^+^ pericytes emerge in parallel with fibrosis progression from *Pdgfrb*-lineage lung pericytes. **(A)** Schematic illustration of lineage-tracing experiments in bleomycin-treated lungs from *Pdgfrb-Cre^ERT2^; ROSA-LSL-YFP* and *αSMA-Cre^ERT2^; ROSA-LSL-RFP* mice. Tamoxifen was administered for 4 days before bleomycin treatment, with lung tissues harvested on day 14. **(B)** Immunostaining of bleomycin-treated lungs from *Pdgfrb-Cre^ERT2^; ROSA-LSL-YFP* (upper) and *αSMA-Cre^ERT2^; ROSA-LSL-RFP* (lower) mice, with Tuj1 (green), lineage-specific reporter protein (gray), and αSMA (red). Arrows indicate Tuj1^+^ pericytes. Scale bars, 8 μm. **(C)** Proportion of YFP- or RFP-expressing cells within Tuj1^+^ pericytes (means ± SEM). **(D)** Immunostaining of bleomycin-treated lungs from *Pdgfrb-Cre^ERT2^; ROSA-LSL-YFP* mouse, with collagen 1 (brown), podoplanin (blue), YFP (gray), and Tuj1 (green). Scale bars, 100 μm. **(E)** Time-course analysis of bleomycin-treated lungs. Upper: collagen 1 (red), Tuj1 (green), Hoechst33342 (blue). Lower: segmentation of Tuj1^+^ pericytes from the upper panels. Scale bars, 80 μm. **(F)** Temporal emergence patterns of Tuj1^+^ pericytes, inflammatory and fibrotic fibroblasts in a publicly available time-course scRNA-seq dataset from the bleomycin model^14^. Relative frequency of these cell types among all cells is calculated for individual mice at specific time points post-injury (*n* = 3). Box plots indicate median (line), interquartile range: IQR (box), 1.5 x IQR (whiskers), and individual values (dots).

### Tuj1^+^ pericytes emerge in parallel with fibrosis progression

To investigate the timing of Tuj1^+^ pericyte emergence during lung fibrogenesis, we analyzed bleomycin-treated lung tissue at multiple time points post-injury (days 0, 3, 5, 7, 10, and 14). Tuj1^+^ pericytes first appeared between days 3 and 5 following bleomycin treatment, with their numbers increasing progressively through day 14, in correlation with the fibrosis progression (Fig. 5 E). To complement these histological observations, we referred to the temporal emergence patterns of inflammatory and fibrotic fibroblasts reported in a recent time-course scRNA-seq study (Tsukui et al., 2024). Inflammatory fibroblasts, characterized by the expression of *Cxcl12*, *Saa3*, *Lcn2,* and interferon-responsive genes, peaked during the inflammatory phase and declined in the fibrotic phase, whereas fibrotic fibroblasts, characterized by the expression of *Cthrc1*, *Col1a1*, and other pathologic extracellular matrix genes, exhibited a progressive increase towards the fibrotic phase (Fig. 5 F). The temporal pattern of Tuj1^+^ pericytes paralleled that of fibrotic fibroblasts, suggesting a potential involvement in the fibrotic process.

### Inflammatory cytokines induce Tuj1 expression in pericytes, while simultaneously upregulating the anti-fibrotic chemokine CXCL10

To investigate the distinctive characteristics of Tuj1^+^ pericytes compared to Tuj1^-^ pericytes, we used the previously analyzed scRNA-seq datasets from the bleomycin model (Tsukui et al., 2020). We initially focused on distinct gene expression profiles between Tuj1^+^ and Tuj1^-^ pericytes. Gene ontology (GO) enrichment analysis of differentially expressed genes in the "activated" pericytes, including Tuj1^+^ pericytes, revealed significant enrichment in categories related to extracellular processes and angiogenesis (Fig. 6 A). These findings prompted us to investigate the secretory properties of Tuj1^+^ pericytes, with a particular focus on their matrisome (Naba et al., 2012) (Fig. 6 B and Fig. S4, A−E). Examination of the matrisome category "secreted factors" revealed a cluster of 12 genes highly expressed in pericytes (Fig. 6 B), identifying *Cxcl10* as the most upregulated secreted factor gene in Tuj1^+^ pericytes. To validate this, we performed RNA *in situ* hybridization for the *Cxcl10* gene in combination with multiplex immunostaining, including Tuj1 (Fig. 6 C). Our results confirmed high *Cxcl10* expression in Tuj1^+^ pericytes associated with PECAM1^+^ capillary ECs within fibrotic regions. Although previous studies have reported CXCL10 secretion from various cell types, including macrophages in the bleomycin model (Tighe et al., 2011) and activated fibroblasts (Tsai et al., 2019), our scRNA-seq analysis revealed that pericytes are among the highest *Cxcl10*-expressing cell types in this fibrosis model (Fig. S4 F). Given that CXCL10 is a well-established angiostatic chemokine (Keane et al., 1997; Keane et al., 1999) and an inhibitor of pulmonary fibrosis (Tager et al., 2004; Jiang et al., 2010), these findings suggest a potential anti-fibrotic role for Tuj1^+^ pericytes in the context of pulmonary fibrosis.

**Figure 6.**
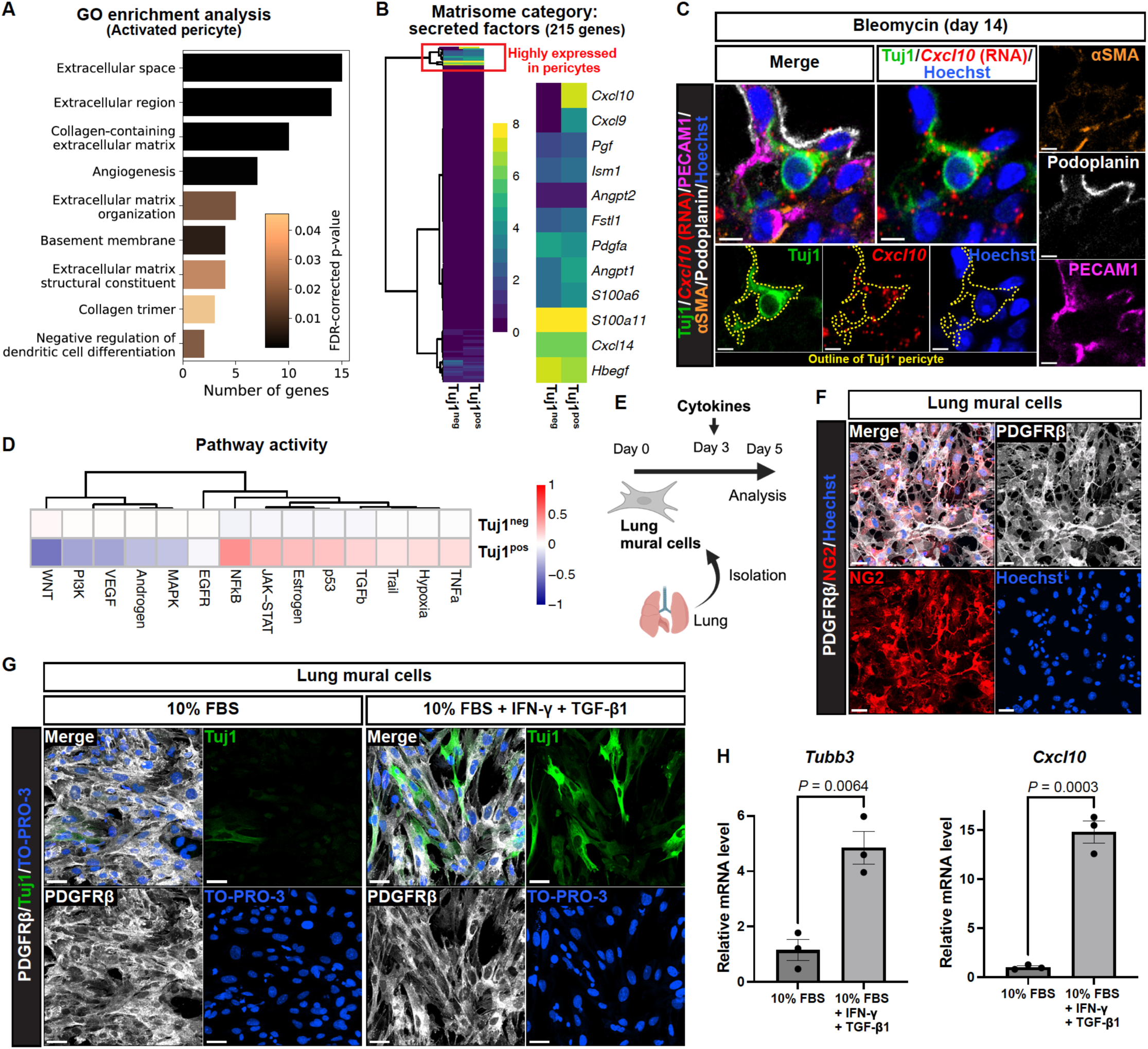
Distinctive characteristics of Tuj1^+^ pericytes compared to Tuj1^-^ pericytes. **(A)** GO enrichment analysis of DEGs upregulated in activated pericytes as analyzed in Fig. 1B. Significantly enriched GO terms are listed with their corresponding FDR-corrected *P* values and associated gene counts. **(B)** Heatmap of average expression levels for 215 secreted factor genes (matrisome category) in Tuj1^neg^ and Tuj1^pos^ pericytes from bleomycin-treated samples in scRNA-seq dataset^21^. A highly expressed gene cluster in pericytes (red rectangle) is shown as an enlarged heatmap with gene names on the right. **(C)** RNA in situ hybridization combined with immunostaining of bleomycin-treated lungs (single z-slice) with *Cxcl10* mRNA (red), Tuj1 (green), PECAM1 (magenta), αSMA (brown), podoplanin (gray), and Hoechst33342 (blue). Scale bars, 5 μm. **(D)** Pathway activity inference of Tuj1^neg^ and Tuj1^pos^ pericytes from bleomycin-treated samples in the scRNA-seq dataset^21^. **(E)** Experimental design to study the effects of inflammatory cytokine on Tuj1 expression using primary lung mural cells. **(F)** Immunostaining of cultured lung mural cells with PDGFRβ (gray), NG2 (red), and Hoechst33342 (blue). Scale bars, 30 μm. **(G)** Immunostaining of cultured lung mural cells with Tuj1 (green), PDGFRβ (gray), and Hoechst33342 (blue). Cultured for 48 hours in growth medium containing 10% fetal bovine serum (FBS), with or without IFN-γ and TGF-β1. Scale bars, 30 μm. **(H)** Relative mRNA expression levels of *Tubb3* and *Cxcl10* in lung mural cells treated with or without IFN-γ and TGF-β1 (*n* = 3 independent experiments, mean ± SEM, unpaired *t* test).

We next investigated mechanisms underlying the emergence of Tuj1^+^ pericytes in the fibrotic regions. Signaling pathway analysis of the previously analyzed scRNA-seq datasets from the bleomycin model (Tsukui et al., 2020) revealed that inflammatory- and fibrosis-associated pathways, including NFκB, JAK-STAT, TGF-β, and TNF-α signaling, were upregulated in Tuj1^+^ pericytes (Fig. 6 D). To test the involvement of inflammatory cytokines for the induction of Tuj1 expression in lung pericytes, we isolated mural cells from mouse lung and cultured them with Interferon-γ (IFN-γ), TNF-α and TGF-β1 (Fig. 6, E and F). Notably, the combination of IFN-γ and TGF-β1 effectively induced Tuj1 expression in lung mural cells in the primary culture (Fig. 4 G and Fig. S4 G). RT-qPCR analysis confirmed that *Tubb3* gene expression was significantly upregulated by this cytokine combination (Fig. 6 H). Furthermore, *Cxcl10* expression was also elevated by this stimulation. Collectively, these data demonstrate that these inflammatory cytokines induce Tuj1 expression in lung pericytes, concomitantly upregulating the anti-fibrotic chemokine, CXCL10.

### Loss of the *Tubb3* gene exacerbates pulmonary fibrosis

To address whether Tuj1 expression in lung pericytes influences the fibrotic process, we analyzed bleomycin-induced pulmonary fibrosis in the mutant mice lacking Tuj1 (*Tubb3*^-/-^ mice). Previous studies have shown that although Tuj1 is widely expressed throughout the central and peripheral nervous systems, *Tubb3^-/-^* mice exhibit no observable neurobehavioral or neuropathological deficits under homeostatic conditions (Latremoliere et al., 2018). Since we analyzed a different line of *Tubb3*^-/-^ mice (Abe et al., 2021), we confirmed that innervation of αSMA^+^ smooth muscle-covered airways and αSMA^+^ VSMC-covered blood vessels in the lungs remained unaffected in our *Tubb3*^-/-^ mice (Fig. 7 A). Our multiplex imaging showed that Tuj1 expression was completely ablated in PDGFRβ^+^/PDGFRα^-^ pericytes in *Tubb3^-/-^* mice, with no changes in the total number of lung pericytes and their Ki67 positivity in the collagen 1^high^ fibrotic regions (Fig. 7, B−D). However, *Tubb3^-/-^*mice exhibited markedly enlarged fibrotic regions, characterized by high expression levels of αSMA and collagen 1 (Fig. 7 E). Quantification of the extent of fibrosis by hydroxyproline assay revealed a significant increase in hydroxyproline content in *Tubb3^-/-^* mice compared to their wild-type littermate controls (Fig. 7 F). Taken together, the absence of Tuj1 expression in lung pericytes appeared to amplify the fibrotic process, suggesting that Tuj1^+^ pericytes may act as negative regulators for fibrosis.

**Figure 7.**
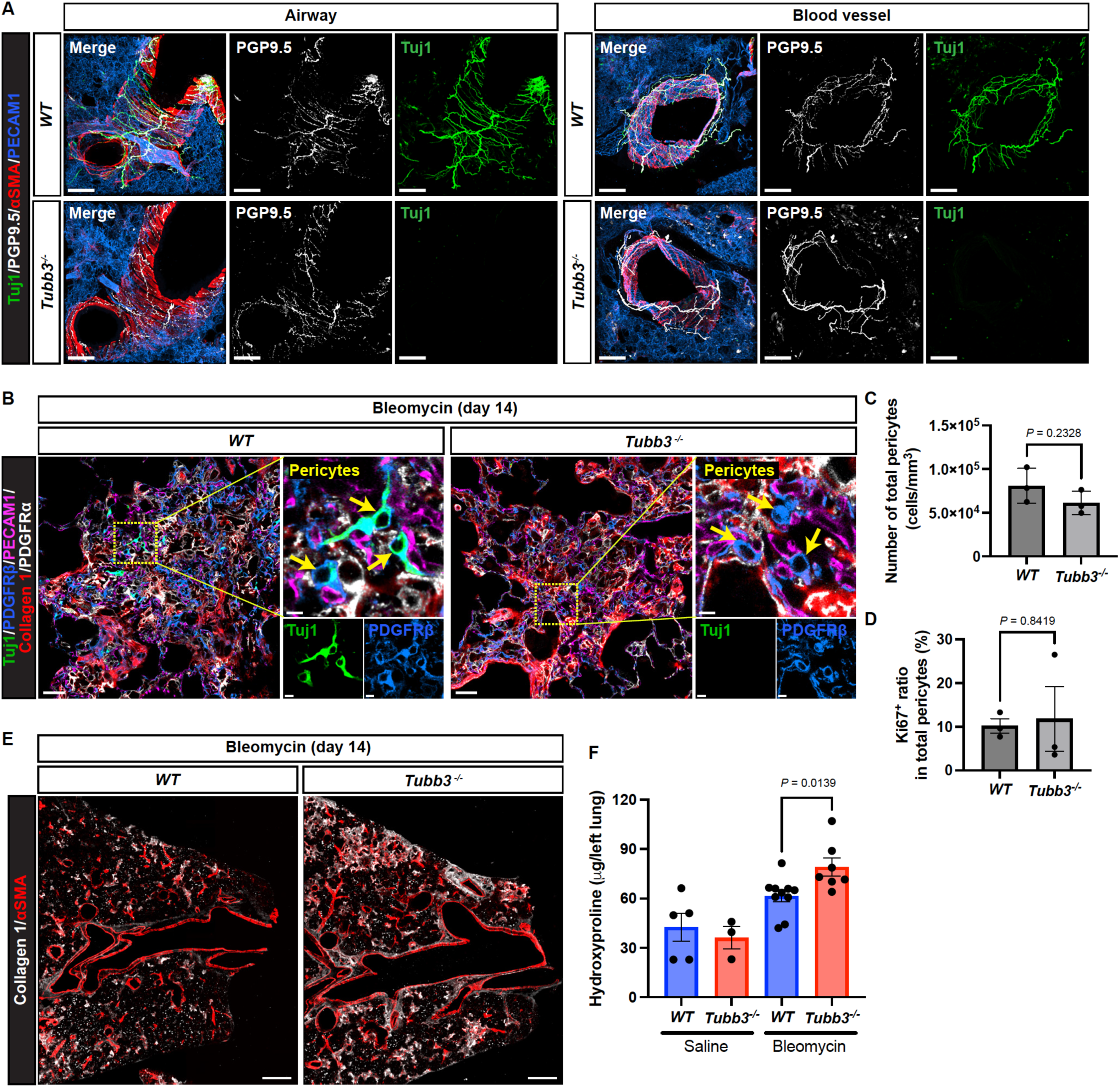
Loss of the Tuj1 gene exacerbates pulmonary fibrosis. **(A)** Immunostaining of lung tissues from wild-type (*WT*) and *Tubb3*^-/-^ mice with antibodies to Tuj1 (green), PGP9.5 (gray), αSMA (red), and PECAM1 (blue). Representative images of airway (left) and blood vessel (right) from *WT* and *Tubb3*^-/-^ mice are shown. Scale bars, 100 μm. **(B)** Immunostaining of bleomycin-treated lungs from *Tubb3*^-/-^ mice and wild-type (*WT*) control littermates (single z-slice) with collagen1 (red), PDGFRα (gray), Tuj1 (green), PDGFRβ (blue), and PECAM1 (magenta). Scale bars, 30 μm (left); 5 μm (magnified views). **(C and D)** Quantification of the total number of pericytes (C) and Ki67^+^ pericytes (D) within the fibrotic regions of *WT* and *Tubb3*^-/-^ mice (*n* = 3 mice per group, mean ± SEM, unpaired *t* test). **(E)** Immunostaining of bleomycin-treated lungs from *WT* and *Tubb3*^-/-^ mice with collagen1 (gray) and αSMA (red). Scale bars, 500 μm. **(F)** Hydroxyproline assay of saline- or bleomycin-treated lung tissues from *WT* and *Tubb3*^-/-^ mice. For *WT* mice, *n* = 5 for saline and *n* = 10 for bleomycin treatment. For *Tubb3*^-/-^ mice, *n* = 3 for saline and *n* = 7 for bleomycin treatment. Mean ± SEM, unpaired *t* test.

### Tuj1^+^ pericytes localize adjacent to pro-fibrotic cells within the fibrotic lung microenvironment

To elucidate the mechanisms underlying the anti-fibrotic role of Tuj1^+^ pericytes, we initially aimed to identify the cellular components within the fibrotic microenvironment that potentially interact with Tuj1^+^ pericytes. Since these pericytes emerge during the fibrotic phase, we focused our spatial imaging analysis on fibroblasts and macrophages, both of which are well-established pro-fibrotic effector cells in fibrosis. Utilizing a 10-color multiplex imaging approach with IBEX (iterative bleaching extends multiplexity) (Radtke et al., 2020), we visualized pro-fibrotic fibroblasts and macrophages, as well as Tuj1^+^ pericytes (Fig. 8 A). Individual cells were then segmented using Cellpose3 (Stringer et al., 2021; Stringer and Pachitariu, 2024) (Fig. 8 B). After quantifying signal intensities across channels and performing K-means clustering on the segmented cells, we annotated Tuj1^+^ pericytes as Tuj1^high^/PDGFRβ^+^/PDGFRα^-^ cells associated with PECAM1^+^ capillary ECs (Fig. 8 C), fibrotic fibroblasts as collagen 1^high^/PDGFRα^+^/PDGFRβ^+^ cells (Fig. 8 D), and pro-fibrotic macrophages as arginase^high^/CD68^+^/CD45^+^ cells (Fig. 8 E). We then conducted spatial analysis of these cell populations using Spatial Patterning Analysis of Cellular Ensembles (SPACE) (Schrom et al., 2024). This analysis successfully delineated non-fibrotic (0-30%) and fibrotic (50-100%) regions along the latent path and detected Tuj1^+^ pericytes in the fibrotic region (Fig. 8 F).

**Figure 8.**
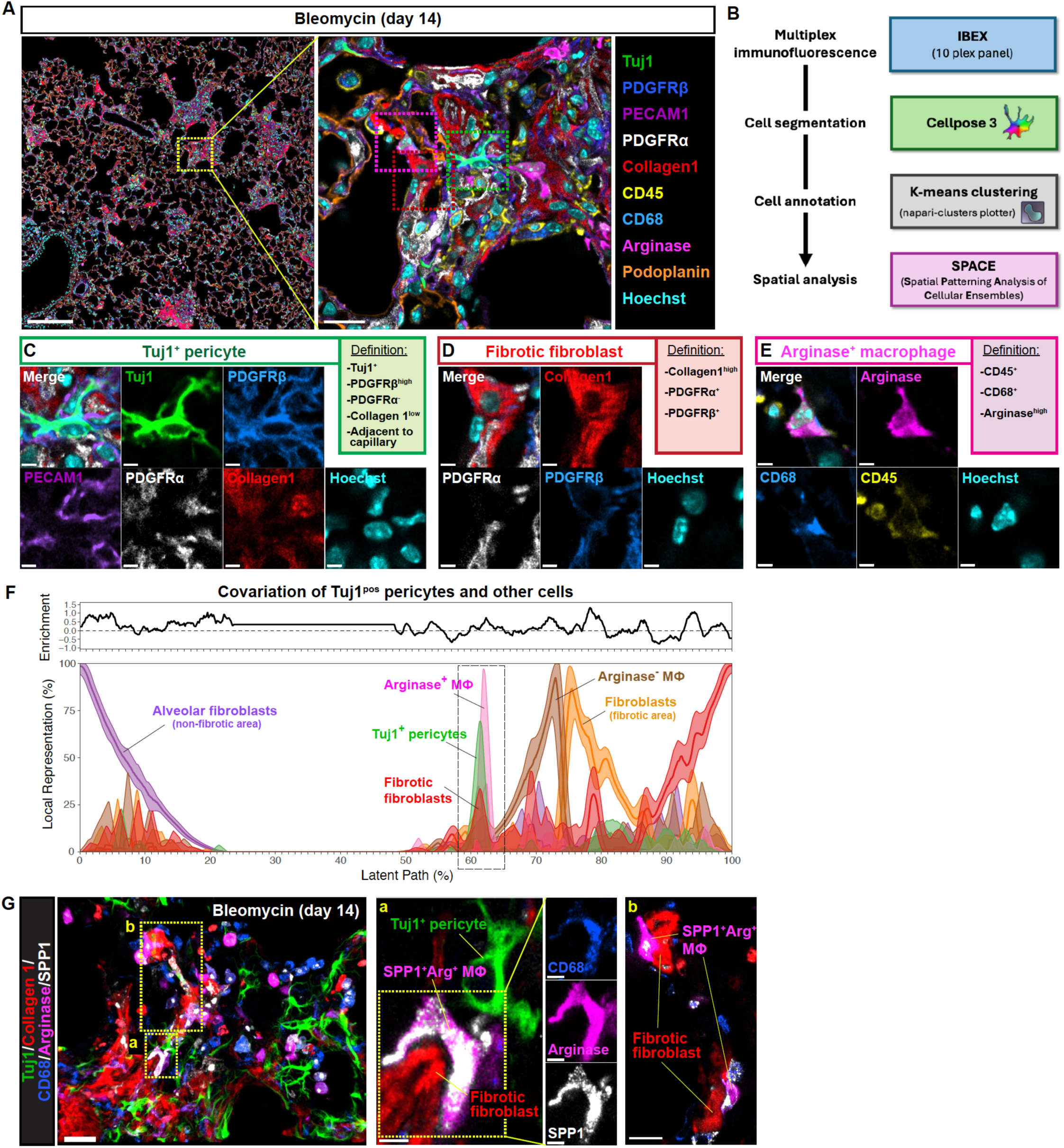
Spatial distribution of Tuj1^+^ pericytes and pro-fibrotic cells in the fibrotic lung microenvironment. **(A)** Immunostaining of bleomycin-treated lungs (single z-slice) with 10 distinct markers using IBEX^34^. Scale bars, 200 μm (left); 20 μm (right). **(B)** Image analysis workflow of the IBEX image. **(C)** Representative image of Tuj1^+^ pericytes, defined as Tuj1^high^/PDGFRβ^+^/PDGFRα^-^ cells associated with PECAM1^+^ capillary ECs. Scale bars, 4 μm. **(D)** Representative image of fibrotic fibroblasts, defined as collagen 1^high^/PDGFRα^+^/PDGFRβ^+^ cells. Scale bars, 4 μm. **(E)** Representative image of arginase^+^ macrophages, defined as arginase^high^/CD68^+^/CD45^+^ cells. Scale bars, 4 μm. **(F)** Spatial analysis of the IBEX image using SPACE^37^. Covariation plots for Tuj1^pos^ pericytes, alveolar fibroblasts (non-fibrotic area), fibroblasts (fibrotic area), fibrotic fibroblasts, arginase^+^ macrophages, and arginase^-^ macrophages. Plots display a smoothed mean and a smoothed 95% confidence interval, along with continuous enrichment score compared to randomized expectations. **(G)** Immunostaining of bleomycin-treated lungs with Tuj1 (green), collagen 1 (red), CD68 (blue), arginase (magenta), and SPP1 (gray). Left: maximum projection image; Right: magnified single-slice images of boxed regions (a, b) in the left panel. Scale bars, 30 μm (maximum projection); 5 μm (A); 15 μm (B). MΦ: macrophage.

Our spatial analysis revealed a positive correlation between the abundance peaks of Tuj1^+^ pericytes and pro-fibrotic macrophages within the fibrotic region (Fig. 8 F; dashed box). In line with this observation, the expression of SPP1 (also known as osteopontin), an additional pro-fibrotic macrophage marker (Morse et al., 2019; Fabre et al., 2023), was also observed in arginase^high^/CD68^+^ pro-fibrotic macrophages in the collagen 1^high^ fibrotic regions (Fig. 8 G; region a). Additionally, an overlapping, albeit less pronounced, peak in fibrotic fibroblasts was observed in the same spatial context (Fig. 8 F; dashed box, and 8 G; region b). These findings suggest potential cell-cell interactions among Tuj1^+^ pericytes, pro-fibrotic macrophages, and fibrotic fibroblasts.

### scRNA-seq analysis revealed the increase of pro-fibrotic cells in *Tubb3*^-/-^ mice

To comprehensively evaluate the impact of Tuj1^+^ pericytes on the neighboring pro-fibrotic cells, we performed scRNA-seq analysis on *Tubb3^-/-^* mice and their wild-type littermate controls. We analyzed whole lung cell samples and lung mural cell-enriched samples from both *Tubb3^-/-^* mice and wild-type controls treated with bleomycin (Fig. 9 A). A total of 25,964 cells were classified into six distinct clusters: fibroblasts, mural cells, immune cells, ECs, epithelial cells, and mesothelial cells (Fig. 9 B). We then conducted cell compositional analysis for each cluster using scCODA (Büttner et al., 2021) to determine which specific cell types within each cluster had cell numbers altered due to the *Tubb3* gene knockout. While the epithelial cell population showed no significant alterations, despite a trend towards increased pre-alveolar type-1 transitional cell state (PATS) cells (Kobayashi et al., 2020) in *Tubb3^-/-^* mice (Fig. S5, A−D), other cell populations exhibited significant changes. In the fibroblast population, *Tubb3^-/-^* mice showed significant increases in fibrotic fibroblasts (e.g., *Cthrc1*^+^, *Runx1*^+^), inflammatory fibroblasts (e.g., *Saa3*^+^, *Sfrp1*^+^, *Lcn2*^+^), and proliferating fibroblasts (*Mki67*^+^) (Fig. 9, C−F). The immune cell population in *Tubb3^-/-^* mice showed significant increases in regulatory T cells (Tregs) and *Spp1*^+^/*Arg1*^+^ macrophages (Fig. 9, G−J and Fig. S5 E). Additionally, the EC population in *Tubb3^-/-^* mice demonstrated significant increases in activated gCap cells (e.g., *Ntrk2* (also known as *TrkB*)^+^, *Lrg1*^+^, *Fxyd5*^+^) along with elevations in proliferating ECs (*Mki67*^+^) and lymphatic ECs (*Prox1*^+^) (Fig. S5, F−I). This pronounced expansion of pro-fibrotic cells in *Tubb3^-/-^* mice corresponds to the observed enhancement of fibrosis in the bleomycin model. Importantly, the increased pro-fibrotic cells, especially fibrotic fibroblasts, *Spp1*^+^/*Arg*1^+^ macrophages, and *Ntrk2*^+^ activated gCap cells, are major neighboring cells of Tuj1^+^ pericytes within lung fibrotic microenvironment, suggesting that Tuj1^+^ pericytes act as negative regulators of the accumulation of these pro-fibrotic cells following bleomycin-induced injury.

**Figure 9.**
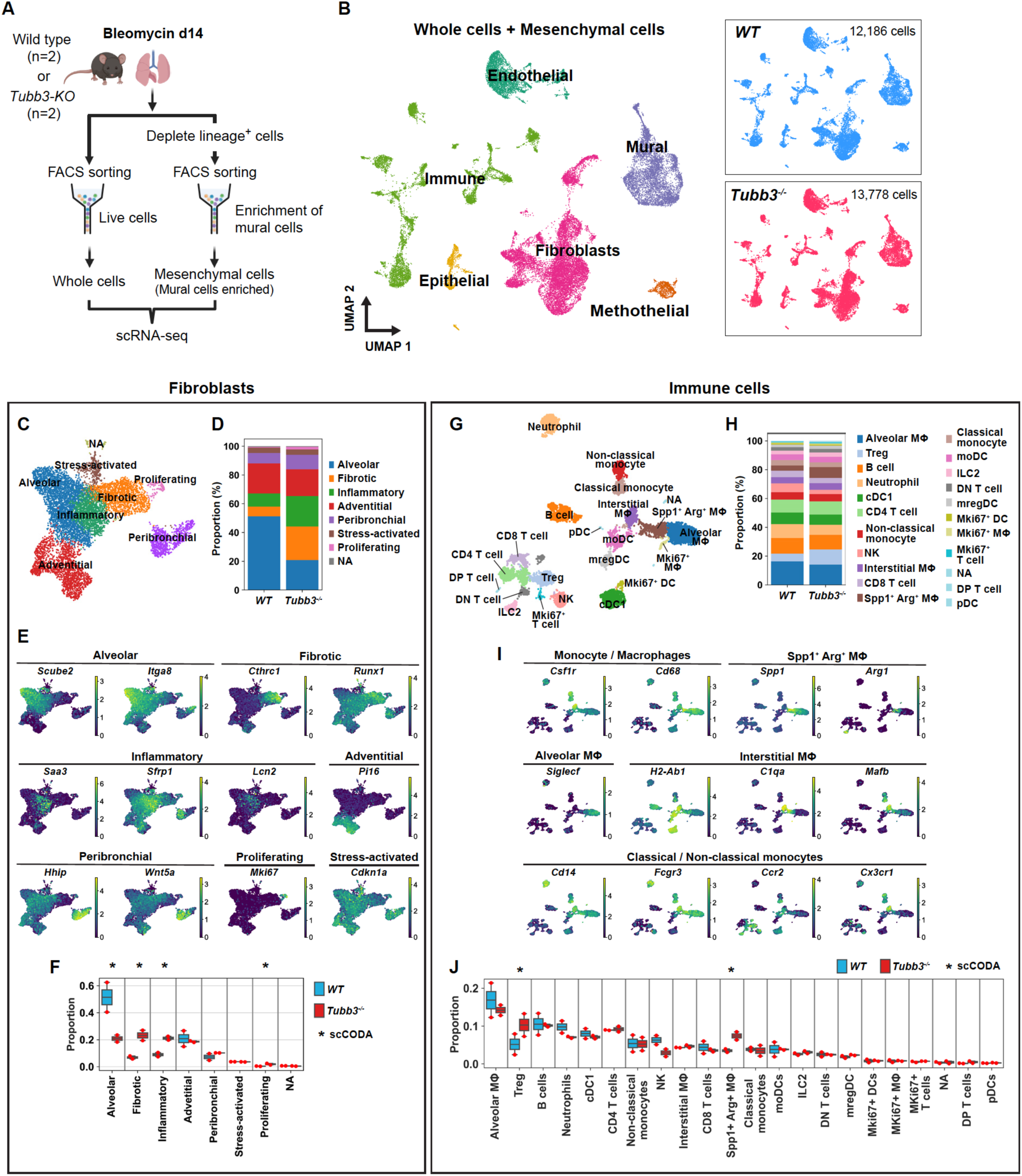
scRNA-seq analysis revealed the increase of pro-fibrotic cells in *Tubb3*^-/-^ mice. **(A)** Schematic illustration of scRNA-seq analysis for bleomycin-treated lung tissues from *WT* and *Tubb3*^-/-^ mice. **(B)** UMAP plots of all cells with cell cluster annotations (left). Separate UMAP plots for *WT* and *Tubb3*^-/-^ samples are displayed on the right panels. **(C−F)** Cell compositional analysis of the fibroblast population from the scRNA-seq data. UMAP plot of the fibroblasts from both *WT* and *Tubb3*^-/-^ mice showing 7 subtypes (C) as defined by marker genes (E). Proportions of each subtype relative to the total fibroblast in *WT* and *Tubb3*^-/-^ mice are shown as stacked barplot (D) and boxplot (F). Statistically credible changes, as tested by scCODA, are noted with an * in panel (F). NA: not applicable. **(G−J)** Cell compositional analysis of the immune cell population from the scRNA-seq data. UMAP plot of the immune cells from both *WT* and *Tubb3*^-/-^ mice showing 21 subtypes (G). Expression of marker genes in monocytes/macrophages is shown in (I). Proportions of each subtype relative to the total immune cells in *WT* and *Tubb3*^-/-^ mice are shown as stacked barplot (H) and boxplot (J). Statistically credible changes, as tested by scCODA, are noted with an * in panel (J). MΦ: macrophage; Treg: regulatory T cell; cDC1: conventional type 1 dendritic cell; NK: natural killer cell; moDC: monocyte-derived dendritic cell; ILC2: type 2 innate lymphoid cell; DN T cell: double negative T cell; mregDC: mature dendritic cell enriched in immunoregulatory molecules; DP T cell: double positive T cell; pDC: plasmacytoid dendritic cell; NA: not applicable.

### CXCL10 suppresses the pro-fibrotic activity of macrophages induced by lung fibroblasts

Given that the aforementioned studies demonstrated preferential expression of the anti-fibrotic and angiostatic chemokine *Cxcl10* in Tuj1^+^ pericytes in the bleomycin model, we next investigated whether the absence of Tuj1 in pericytes affects *Cxcl10* expression. To address this, we first utilized our scRNA-seq datasets to examine *Cxcl10* expression in pericytes. From the mural cell population in the datasets derived from *Tubb3^-/-^*mice and wild-type controls (Fig. 10 A), we subset the pericyte population (Fig. 10 B). Consistent with the aforementioned multiplex immunostaining results (Fig. 7, B−D), the total number of pericytes in the lung remained comparable between *Tubb3^-/-^* mice and wild-type controls (Fig. 10 C). We subsequently analyzed transcriptional changes in pericytes resulting from *Tubb3* gene knockout. As expected, *Tubb3* expression was significantly reduced in *Tubb3^-/-^* mice (Fig. 10 D). Intriguingly, despite this reduction, we observed upregulation of activated pericyte markers, including *Rgs5* and *Col18a1*, suggesting enhanced pericyte activation in *Tubb3^-/-^* mice. In contrast, *Cxcl10* expression tended to be downregulated in the pericytes of *Tubb3^-/-^* mice (Fig. 10 D). To validate this finding, we conducted RNA *in situ* hybridization for the *Cxcl10* gene, combined with multiplex immunostaining to visualize PDGFRβ^+^/PDGFRα^-^ pericytes associated with PECAM1^+^ capillary ECs in the collagen 1^high^ fibrotic regions (Fig. 10 E). We quantified the *Cxcl10* spot signals in pericytes (Fig. 10 F). Consistent with the scRNA-seq data, we observed a significant reduction in *Cxcl10* expression in the pericytes of *Tubb3^-/-^* mice. This reduction in the anti-fibrotic *Cxcl10* in pericytes may contribute to the enhancement of fibrosis observed in these mutant mice.

**Figure 10.**
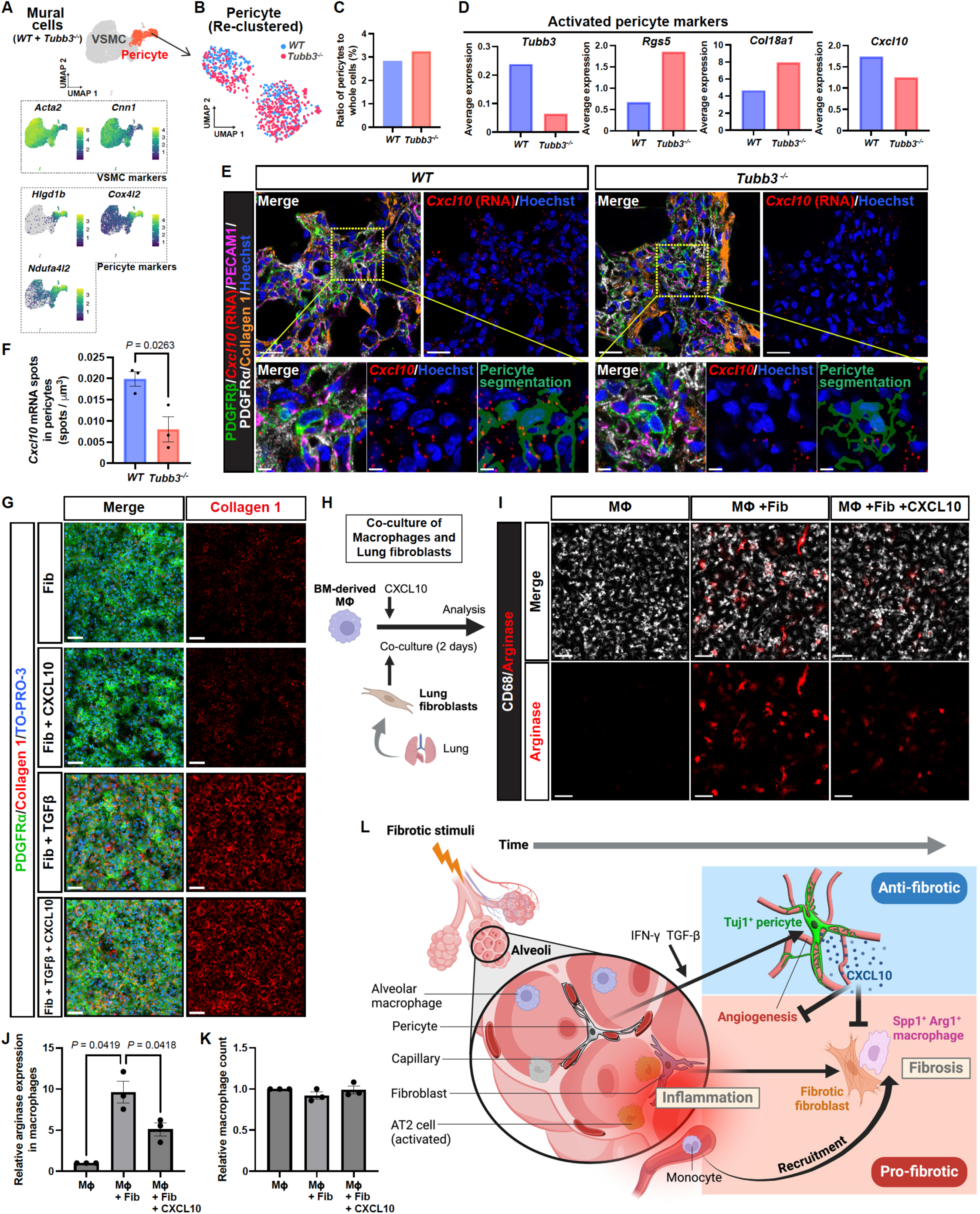
CXCL10 suppresses the pro-fibrotic activity of macrophages induced by lung fibroblasts. **(A)** UMAP plots showing expression of marker genes for mural cell subtypes (VSMC and pericyte) from the scRNA-seq analysis of *WT* and *Tubb3*^-/-^ mice. **(B−D)** scRNA-seq analysis of the pericyte population. **(B)** UMAP plot of re-clustered *WT* pericyte (blue) and *Tubb3*^-/-^ pericyte (red) population. **(C)** Percentage of *WT* and *Tubb3*^-/-^ pericytes relative to total lung cells. **(D)** Average expression levels of activated pericyte marker genes (*Tubb3*, *Rgs5*, and *Col18a1*) and *Cxcl10* in *WT* and *Tubb3*^-/-^ pericytes. **(E)** RNA in situ hybridization combined with immunostaining of bleomycin-treated lungs from *WT* and *Tubb3*^-/-^ mice (single z-slice) with *Cxcl10* mRNA (red), PDGFRβ (green), PDGFRα (gray), PECAM1 (magenta), collagen 1 (brown), and Hoechst33342 (blue). Pericyte segmentation was performed based on the PDGFRβ, PDGFRα and PECAM1 staining patterns. Scale bars, 20 μm (upper); 5 μm (lower). **(F)** Quantification of *Cxcl10* RNA spots within the segmented pericyte areas in fibrotic regions of *WT* and *Tubb3*^-/-^ mice. (*n* = 3 mice, mean ± SEM, unpaired *t* test). **(G)** Immunofluorescence images of lung fibroblasts labeled with PDGFRα (green), collagen 1 (red), and the nuclear marker TO-PRO-3 (blue). The freshly isolated lung fibroblasts were cultured for 48 hours in growth medium containing 10% fetal bovine serum (FBS), either alone or supplemented with CXCL10 or TGF-β1, or TGF-β1 and CXCL10 together. Fib: lung fibroblasts; TGFβ: TGF-β1. Scale bars, 50 μm. **(H)** Experimental design to study the effect of CXCL10 on fibroblast-mediated pro-fibrotic differentiation of bone marrow (BM)-derived macrophages in culture. **(I)** Immunostaining of cultured BM-macrophages alone, BM-macrophages co-cultured with lung fibroblasts, and BM-macrophages co-cultured with lung fibroblasts in the presence of CXCL10, labeled with CD68 (gray) and arginase (red). Scale bars, 50 μm. MΦ: BM-derived macrophages; Fib: lung fibroblasts. **(J and K)** Quantification of arginase expression (J) and total number of macrophages (K). Arginase intensity values from three independent experiments were normalized to their respective macrophage-only controls and combined for statistical analysis (*n* = 3 independent experiments, mean ± SEM, one-way ANOVA followed by Tukey’s multiple comparisons test). **(L)** Schematic model illustrating the anti-fibrotic role of Tuj1^+^ lung pericytes during pulmonary fibrosis. Following fibrotic stimuli, damaged type II alveolar epithelial cells (AT2 cells) trigger local inflammation, leading to the recruitment and differentiation of circulating monocytes into pro-fibrotic macrophages (Spp1^+^/Arg1^+^). Concurrently, activated alveolar fibroblasts at inflammatory sites differentiate into fibrotic fibroblasts (collagen 1^high^). In parallel with these pro-fibrotic responses, the local inflammation also activates lung pericytes through IFN-γ and TGF-β signaling, inducing Tuj1 expression. These Tuj1^+^ pericytes upregulate the anti-fibrotic chemokine CXCL10, which suppresses both angiogenesis and pro-fibrotic activity in macrophages, thereby counteracting the progression of pulmonary fibrosis.

The marked increase in fibrotic fibroblasts and activated gCap ECs in *Tubb3^-/-^* mice could be explained by the decreased expression of the anti-fibrotic and angiostatic chemokine *Cxcl10* in pericytes. Given previous studies showing minimal effects of CXCL10 on fibroblast proliferation and collagen production in culture (Tager et al., 2004; Jiang et al., 2010), it is possible that CXCL10 indirectly influences fibrotic fibroblasts. To address this, we cultured freshly isolated lung fibroblasts and examined whether CXCL10 could inhibit their induction of collagen 1 by TGF-β1. Consistent with the previous studies, we found no significant effect of CXCL10 on collagen 1 expression in these activated fibroblasts (Fig. 10 G), suggesting that the increased number of fibrotic fibroblasts in *Tubb3^-/-^*mice may be a secondary effect resulting from the reduction of *Cxcl10* in pericytes.

Given the significant role of pro-fibrotic macrophages in the fibrotic process, we hypothesized that CXCL10 may influence the pro-fibrotic activity of these cells. To address this, we developed a co-culture system of CD68^+^ bone marrow (BM)-derived macrophages and freshly isolated lung fibroblasts to induce arginase^+^/CD68^+^ pro-fibrotic macrophages (Fig. 10, H and I). Interestingly, CXCL10 inhibited the arginase induction in macrophages in the co-culture, without affecting the total number of macrophages (Fig. 10, I−K). These findings indicate that CXCL10 plays a suppressive role in pro-fibrotic macrophage differentiation. Taken together, these findings suggest that the increased fibrosis observed in *Tubb3^-/-^* mice may be largely due to the increased number of pro-fibrotic macrophages, driven by the reduction of *Cxcl10* in pericytes.

## Discussion

Here, we demonstrate a novel anti-fibrotic state of pericytes in pulmonary fibrosis. Our findings suggest that in response to fibrotic stimuli, concurrent with the emergence of pro-fibrotic fibroblasts and macrophages, lung pericytes express Tuj1, adopting an anti-fibrotic phenotype. This phenotype acts as a negative regulator of pro-fibrotic macrophage differentiation through the release of CXCL10, thereby counteracting the progression of pulmonary fibrosis (Fig. 10 L).

Activated pericytes are known to proliferate and associate with activated capillary ECs within fibrotic regions. However, Tuj1 expression does not appear to influence pericyte proliferation, as observed in the lack of difference in pericyte proliferation between Tuj1^+^ and Tuj1^-^ pericytes within fibrotic regions, as well as the absence of changes in total pericyte numbers and their proliferative state in *Tubb3^-/-^*mice. Notably, in response to fibrotic stimuli, activated pericyte markers are further upregulated in *Tubb3*^-/-^ mice. Since most activated pericytes within fibrotic regions express Tuj1, and the absence of Tuj1 in these pericytes exacerbates fibrosis, this expression may contribute to an anti-fibrotic state in pericytes during pulmonary fibrosis. Although pericytes have long been considered a source of myofibroblasts, our findings advance this emerging paradigm by demonstrating an anti-fibrotic role for Tuj1^+^ activated pericytes in pulmonary fibrosis, highlighting the importance of pericytes as a counteracting cellular component in mitigating fibrotic processes.

Tuj1 expression is observed in both pericytes and VSMCs in a bleomycin-induced pulmonary fibrosis model, although pericytes constitute most Tuj1-expressing cells within fibrotic regions. To directly confirm that Tuj1 expression in pericytes is necessary for inducing anti-fibrotic effects in pulmonary fibrosis, pericyte- or VSMC-specific deletion of the *Tubb3* gene would be required. Given the historical challenge of identifying highly specific markers and genetic *Cre* driver mice for pericytes (Yao, 2022), the recent discovery of *Higd1b* gene as a relatively unique marker for lung pericytes has enabled the generation of inducible *Higd1b-CreER* driver mice (Klouda et al., 2025). These mice could serve as a valuable tool for pericyte-specific deletion of the conditional *Tubb3* allele. Alternatively, genetically ablating Tuj1-expressing cells could validate their anti-fibrotic effect within fibrotic regions. Since Tuj1 is constitutively expressed in all neurons, targeted ablation of Tuj1^+^ pericytes would be needed to assess their anti-fibrotic effects in pulmonary fibrosis.

How do Tuj1^+^ pericytes contribute to an anti-fibrotic state during pulmonary fibrosis? One possibility is that Tuj1^+^ pericytes release anti-fibrotic factors that suppress pro-fibrotic cells, including activated capillary ECs, pro-fibrotic fibroblasts, and macrophages. Notably, Tuj1^+^ pericytes express the anti-fibrotic chemokine *Cxcl10,* and the absence of Tuj1 reduces *Cxcl10* expression in pericytes, accompanied by the marked increase in the aforementioned pro-fibrotic cells. The in vitro data demonstrate that CXCL10 does not affect collagen production by fibrotic fibroblasts; rather, it suppresses the pro-fibrotic differentiation of BM-derived macrophages in the co-culture with primary lung fibroblasts. Recent studies reveal that arginase1 (Arg1)^+^ pro-fibrotic macrophages are found in close proximity to pro-fibrotic fibroblasts within fibrotic regions, and that macrophage-specific deletion of *Arg1* attenuates pulmonary fibrosis following fibrotic stimuli, suggesting that Arg1^+^ pro-fibrotic macrophages play an essential role in pulmonary fibrosis (Yadav et al., 2024). Moreover, fibroblast-derived IL-6 has been shown to induce Arg1 expression in macrophages in culture (Yadav et al., 2024). In our studies, Tuj1^+^ pericytes are also localized in close proximity to Arg1^+^ pro-fibrotic macrophages within fibrotic regions, and the absence of Tuj1 in pericytes exacerbates pulmonary fibrosis following fibrotic stimuli, suggesting that Tuj1^+^ pericytes negatively regulate these macrophages through CXCL10. However, whether CXCL10 inhibits the cross-talk between macrophages and fibroblasts or directly suppresses the pro-fibrotic differentiation of macrophages remains to be determined.

Although Tuj1 expression is predominantly specific to the nervous system, its expression in non-neuronal human tumors has been reported to be associated with resistance to tubulin-binding agents, high metastatic potential, and poor patient outcomes (Kanakkanthara and Miller, 2021). Various signaling pathways, including nuclear hormone receptor signaling, bromodomain extraterminal domain (BET) proteins-mediated epigenetic regulation, hypoxia-mediated signaling, and microRNAs, have been reported to regulate Tuj1 expression in cancer cells (Kanakkanthara and Miller, 2021), suggesting that multiple regulatory mechanisms influence Tuj1 expression across diverse cancer types. In our studies, Tuj1 expression displays a strong correlation with CXCL10 (also known as IFN-γ-induced protein 10), potentially linking Tuj1 expression in pericytes to molecular alterations associated with inflammatory signaling. However, the absence of Tuj1 in pericytes leads to the marked reduction in *Cxcl10* expression. These findings suggest that Tuj1-mediated signaling pathways may regulate *Cxcl10* expression. Further investigation is required to understand the link between Tuj1-mediated signaling and *Cxcl10* expression in activated pericytes in pulmonary fibrosis. This will provide therapeutic strategies to enhance the anti-fibrotic properties of lung pericytes, specifically by increasing CXCL10 expression, aiming to treat pulmonary fibrosis.

## Materials and methods

### Mice

All animal procedures were approved by the National Heart, Lung, and Blood Institute (NHLBI) Animal Care and Use Committee in accordance with NIH research guidelines for the care and use of laboratory animals. The following mice were used in this study: *Pdgfrb-Cre^ERT2^* mice (The Jackson Laboratory, 029684), *R26R-EYFP* mice (The Jackson Laboratory, 006148), *αSMA-Cre^ERT2^* mice (Wendling et al., 2009), *NG2-Cre^ERT2^* mice (The Jackson Laboratory, 0068538), *Scube2-Cre^ERT2^*mice (Tsukui et al., 2024), *Cthrc1-Cre^ERT2^* mice (Tsukui et al., 2024), *R26R-tdTomato* mice (The Jackson Laboratory, 007914), and *Col-GFP* mice (Yata et al., 2003). The *Cre*-mediated excision was induced by administrating tamoxifen via intraperitoneal injection (I.P.). As *Tubb3^-/-^* mice, *Tubb3*-*Cre* knockin mice in which the *Cre* transgene was inserted in-frame at the translation initiation site of the *Tubb3* gene (Abe et al., 2021) were used. Homozygous *Tubb3^-/-^* mice were obtained by crossing heterozygous *Tubb3*-*Cre* knockin mice. Mice between the ages of 8 and 11 weeks old were used for the experiments. Male mice were used for the hydroxyproline experiment. Both male and female mice were used in the other experiments. For fibrosis induction, a dose of 0.8 U/kg bleomycin was intratracheally administered into mice.

### Lung tissue preparation for imaging analysis

Lung tissues were inflated with 4% paraformaldehyde (PFA) through the trachea and removed from mice. The tissues were then washed with phosphate-buffered saline (PBS) and fixed with 4% PFA or BD Cytofix (BD Biosciences, BDB554655) diluted in PBS (1:3) overnight at 4°C. After washing in PBS, the fixed tissues were incubated in 30% sucrose/PBS overnight at 4°C, embedded in Tissue-Tek O.C.T. Compound (Sakura), and stored at −80°C.

### Multiplex immunofluorescence of lung tissue

Cryosections (30-100 µm) from fixed-frozen tissue embedded in O.C.T. Compound were adhered to Superfrost Plus Gold slides (Fisher Scientific, 15-188-48) or two-well Chambered Coverglasses (Lab-Tek, 155379) coated with chrome alum gelatin (Newcomer Supply, Part 1033A). Sections were washed with PBS and PBS-T (0.1% Tween 20), permeabilized with 0.5% Triton X-100 in PBS for 30 minutes, and then blocked with either 1% bovine serum albumin (BSA; Millipore Sigma, A9418) in PBS with 0.1% Tween 20 or 10% donkey serum in PBS with 0.1% Tween 20 for 1 hour at room temperature. Primary and secondary antibodies used in this study are listed in Table S1. Primary antibodies were incubated in blocking buffer overnight at 4°C. Fluorescence-conjugated secondary antibodies were used at a 1:250 dilution in a blocking buffer (with Hoechst 33342 (Biotium, 40046, 1:500) when needed) and incubated for 2-5 hours at room temperature or overnight at 4°C. After washing in PBS-T, sections were mounted with ProLong Gold Antifade Mountant (Thermo Fisher Scientific, P36934), Ce3D+ (Anderson et al., 2020) for tissue clearing, or Fluoromount-G (Southern Biotech, 0100-01) for iterative bleaching extends multiplexity (IBEX) (Radtke et al., 2020).

Cyclic immunofluorescence was performed using the IBEX protocol. After confocal imaging, mounting media was thoroughly removed by washing with PBS. The sections were then incubated with lithium borohydride (LiBH4, 1 mg/mL; Stem Chemicals, 16949-15-8) for 30 minutes under white light to bleach fluorescent signals. After washing in PBS, subsequent cycles were started from the primary antibody reaction. Hoechst 33342 was used as the fiducial marker for image registration between cycles.

### Human lung tissue

Fibrotic lung tissue samples were obtained from lung cancer patients with IPF during surgical resection at Kumamoto University Hospital. This study was approved by the Institutional Review Board of Kumamoto University Hospital (approval number: 2460). The lung tissues were fixed in 4% PFA and subjected to hematoxylin and eosin (H&E) staining or multiplex immunofluorescence analysis.

### RNA in situ hybridization for *Col1a1* combined with immunofluorescence

RNA in situ hybridization was performed using HCR RNA-FISH technology (Molecular Instruments). Cryosections (30-50 µm) from fixed-frozen tissue embedded in O.C.T. Compound were adhered to Superfrost Plus Gold slides coated with chrome alum gelatin. Target retrieval was conducted by incubating sections with 0.5% Triton X-100 in PBS for 10 minutes at room temperature. In situ hybridization with the *Col1a1* probe set, followed by amplification using an Alexa 647-conjugated B3 amplifier were carried out, in accordance with the manufacturer’s instructions. Following the hybridization, immunofluorescence staining was performed. Sections were washed in PBS-T (0.2% Triton X-100) and blocked with 10% goat serum in PBS containing 0.2% Triton X-100 for 1 hour at room temperature. Primary antibodies, including an Alexa 488-conjugated anti-Tuj1 antibody (Biolegend, 801203; 1:200) and a Cy3-conjugated anti-αSMA antibody (Millipore Sigma, C6198; 1:500), were incubated in the blocking buffer overnight at 4°C. After washing in PBS-T, sections were mounted using ProLong Gold Antifade Mountant.

### RNA in situ hybridization for *Cxcl10* combined with IBEX

To detect *Cxcl10* RNA, which has limited target sites for probe binding due to its small size, we employed Yn situ (Wu et al., 2022), a sensitive RNA detection method. This method was integrated with IBEX to simultaneously detect *Cxcl10* RNA and protein signals. The buffer composition for RNA in situ hybridization was adapted from Murakami et al. (Murakami and Heintz, 2022).

Cryosections (30 µm) of fixed-frozen tissue embedded in O.C.T. Compound were adhered to Superfrost Plus Gold slides coated with chrome alum gelatin. The sections were post-fixed with 4% PFA for 10 minutes at room temperature. After washing with PBS, the sections were incubated in Methylimidazole buffer for 5 minutes, followed by 1-ethyl-3-(3-dimethylaminopropyl) carbodiimide (EDC) fixative for 1 hour at room temperature, in accordance with the protocol by Wu, et al. (Wu et al., 2022). After another PBS wash, the sections underwent dehydration through a graded methanol series (30%, 50%, 80%, and 100%, twice, 5 minutes each). The slides were then stored in 100% methanol at −80°C until use. Prior to use, the slides were brought to room temperature and rehydrated through a decreasing methanol gradient (80%, 50%, 30%, 5 minutes each).

To detect protein combination 1 (Tuj1, PECAM1, αSMA, and podoplanin), rehydrated sections were treated with 10 mM HCl for 30 minutes at room temperature. Following a 10-minute wash with washing buffer 1 (5×SSC, 0.01% Tween 20, and 20 mM citric acid; w.b.1), the sections were incubated with washing buffer 2 (2×SSC, 0.01% Tween 20, and 4 mM citric acid; w.b.2) containing 0.1 µg/mL proteinase K (Thermo Fisher Scientific, 25530049) for 10 minutes at 37°C. For the detection of protein combination 2 (PDGFRβ, PECAM1, PDGFRα, and collagen 1), rehydrated sections were washed with w.b.1 for 10 minutes and then incubated with w.b.2 containing 10 µg/mL proteinase K for 10 minutes at 37°C. Following the proteinase K treatment, the sections were washed sequentially with w.b.1 (5 minutes), PBS (15 minutes), and PBS-T (0.1% Tween 20, 15 min). The sections were blocked with 1% BSA in PBS containing 0.1% Tween 20 for 1 hour at room temperature. Primary and secondary antibodies are listed in Table S1. Primary antibodies were incubated in the blocking buffer overnight at 4°C. Fluorescence-conjugated secondary antibodies (1:250) and Hoechst 33342 (1:500) were applied in the blocking buffer and incubated overnight at 4°C. After washing with PBS-T, the sections were mounted with Fluoromount-G for confocal scanning. Following confocal imaging, the mounting medium was thoroughly removed with PBS washing. Fluorescent signals were bleached by incubating the sections with LiBH4 (1 mg/mL) for 30 minutes under white light.

After PBS washing, Yn situ was performed. The sections were washed with w.b.1 for 10 minutes at room temperature and incubated with w.b.2 containing 10 µg/mL proteinase K for 30 minutes at 37°C. After w.b.1 washing (5 minutes), the sections were pretreated with hybridization buffer (5×SSC, 0.01% Tween 20, 30% formamide, and 3% PEG8000) for 5 minutes at room temperature. The sections were then incubated with the *Cxcl10* probe set (4 pmol per probe, listed in Table S2), designed using insitu_probe_generator (Kuehn et al., 2022), in hybridization buffer overnight at 37°C. Following four washes with w.b.1 (30 minutes each, 37°C), the sections were pretreated with hybridization buffer for 5 minutes at room temperature. The sections were incubated with a preamplifier (15 ng/µL), synthesized from plasmid A1-20B1 (P004) (Addgene, #184056) using a nickase Nt.BspQI (New England Biolabs, R0644S), in hybridization buffer for 3 hours at 37°C. After four washes with w.b.1 (15 min each, 37°C), a thermocycler-annealed Alexa 647-conjugated B1 amplifier (3 µM; Molecular Instruments) was diluted in hybridization buffer (1:50) and applied overnight at room temperature. Following four washes with w.b.1 (15 minutes each, 37°C), the sections were incubated with w.b.2 containing Hoechst 33342 (1:500) for 15 minutes at 37°C. After w.b.2 washing (15 minutes, 37°C), the slides were mounted with ProLong Gold Antifade Mountant. Hoechst 33342 was used as the fiducial marker for image registration between cycles.

### Image Analysis

Confocal microscopy was performed using Leica TCS SP5 or SP8 confocal microscopes (Leica). For publication-quality images, brightness/contrast adjustments were applied uniformly to all comparable images. Unless otherwise stated, images are presented as maximum intensity projections (MIP) of tiled z-stacks using Fiji (Schindelin et al., 2012), Imaris (Oxford Instruments), or napari (napari contributors, 2019). Image alignment of all IBEX panels was performed using SITK-IBEX (Radtke et al., 2020) with Hoechst staining serving as the fiducial marker.

Quantification of Tuj1^+^ pericytes or Tuj1^+^ VSMCs in Figs. 4C, 5C, S3B was performed using the napari-3d-counter plugin.

In Figs. 2B, 2C, 2G, and 5E, area annotation and Tuj1^+^ cell segmentation were performed using the Fiji plugin Labkit (Arzt et al., 2022). In Fig. 2C, the density of Tuj1^+^ cells was calculated using Gaussian kernel density estimation and visualized as a heatmap overlaid on the area annotation image.

In Figs. 2E, 2F, 2H, 3B, 3C, 7C, and 7D, four distinct pericyte populations (Tuj1^+^ Ki67^+^, Tuj1^+^ Ki67^-^, Tuj1^-^ Ki67^+^, and Tuj1^-^ Ki67^-^) were counted using the Fiji plugin MoBIE (Pape et al., 2023). Individual channel images and instance-segmented nuclei labels (generated using Cellpose3 (Stringer et al., 2021; Stringer and Pachitariu, 2024)) were imported into a MoBIE project, where the four distinct pericyte populations were annotated to their corresponding nuclei objects.

In Figs. 1J and S2C, 3D cropping was performed using the Fiji plugin 3Dscript (Schmid et al., 2019).

In Fig. 6F, whole-cell segmentation in the IBEX 10 multiplex image utilized PDGFRα, PDGFRβ, PECAM1, podoplanin, and collagen 1 channels as cytoplasm markers, with Hoechst as the nuclear marker. Enhance Contrast and Difference-of-Gaussian filter in 3D were applied to each channel image. For the cytoplasm markers, each channel was subtracted from all other channels, and the resulting images were merged to create a composite cytoplasm image. The processed cytoplasm and Hoechst images were input into Cellpose3 for 3D cell segmentation using the cyto3 model following image restoration. After segmentation, imaging artifacts were removed through size filtering and exclusion of cell labels with low Hoechst intensity. Cell clustering was performed using UMAP dimensionality reduction, followed by K-means clustering (k=25) on standardized mean intensity values from raw images. The resulting clusters were then manually annotated using the napari-clusters-plotter plugin. The annotated cell data was then analyzed using SPACE (Schrom et al., 2024), with results visualized using SPACE’s built-in visualization functions.

In Fig. 10F, *Cxcl10* RNA spots were quantified using U-FISH (Xu et al., 2024). Pericyte segmentation was performed using Labkit. For the pericyte segmentation, PDGFRβ, PDGFRα, PECAM1, and collagen 1 channel images were pre-processed with Enhance Contrast and Difference-of-Gaussian filter in 3D. The PDGFRβ image was then subtracted from all other processed channels, and the resulting PDGFRβ images were used for the pericyte segmentation using Labkit. *Cxcl10* RNA spots were quantified within these segmented pericyte areas by U-FISH.

In Figs. 10J and 10K, tiled z-stack images were processed as maximum intensity projections. Macrophages were segmented using Cellpose3 with the CD68 channel as the cytoplasmic marker. Following size-based filtering to remove imaging artifacts, arginase intensity from raw images was measured for each segmented macrophage.

### Hydroxyproline assay

Pulmonary fibrosis after bleomycin treatment was assessed by measuring hydroxyproline content in the left lobes of mouse lung tissue using a Hydroxyproline Assay Kit (Cell Biolabs Inc., STA-675) in accordance with the manufacturer’s protocol.

### Lung tissue dissociation

Mouse lungs were harvested following perfusion with 5 mL of PBS through the right ventricle. The tissue was mechanically dissociated using scissors and digested in an enzyme solution containing 0.25% Collagenase A (Millipore Sigma, 10103586001), 1 U/mL Dispase II (Millipore Sigma, 4942078001), and 2,000 U/mL DNase I (Millipore Sigma, 260913-10MU) in Hanks’ Balanced Salt Solution. The tissue suspension was incubated at 37°C for 60 minutes with gentle trituration using a micropipette every 20 minutes. Following digestion, the cell suspension was filtered through a 100-μm cell strainer (BD Biosciences), washed with PBS, and resuspended in PBS containing 0.5% BSA.

### In vitro stimulation of primary lung mural cells

Lung mural cells were isolated by magnetic negative selection using the following antibodies: rat anti-mouse CD31 (BD Biosciences, 553370; 1:100), CD45 (eBioscience, 14-0451-85; 1:100), EpCAM (Biolegend, 118201; 1:100), and TER-119 (eBioscience, 14-5921-85; 1:100) antibodies, followed by the addition of sheep anti-rat Dynabeads (Thermo Fisher Scientific, 11035). Subsequently, an additional negative selection was performed using anti-mouse PDGFRα MicroBeads (Miltenyi Biotec, 130-101-502; 1:50). Following magnetic separation, 1 × 10^5^ cells were seeded into collagen type I-coated glass-bottom dishes with cloning rings and cultured in basal medium (McErlain et al., 2024) supplemented with 10% fetal bovine serum (FBS; Hyclone, SH3007) and 1% penicillin-streptomycin (Gibco, 15140122) for 72 hours. The medium was then replaced with fresh medium containing cytokines in the following combinations: IFN-γ (50 ng/mL; PeproTech, 315-05), TGF-β1 (1 ng/mL, PeproTech, 100-21), or TNF-α (10 ng/mL; PeproTech, 315-01A) alone; IFN-γ + TGF-β1, IFN-γ + TNF-α, or TGF-β1 + TNF-α in combination; or all three cytokines (IFN-γ + TGF-β1 + TNF-α) together. The cells were incubated with these cytokine treatments for 48 hours. Following cytokine stimulation, the cells were fixed with 4% PFA in PBS and processed for immunostaining. For RNA isolation, total cells isolated from a single mouse were divided into two and seeded into collagen type I-coated 96-well plates and cultured as described above. After cytokine stimulation, the cells were lysed using RLT buffer from the RNeasy Plus Micro Kit (Qiagen, 74034) and processed for RNA isolation. All cell culture procedures were performed under standard conditions (37°C, 5% CO_2_).

### In vitro stimulation of primary lung fibroblasts

Lung fibroblasts were isolated by magnetic negative selection using rat anti-mouse CD31 (1:100), CD45 (1:100), EpCAM (1:100), TER-119 (1:100), and MCAM (Biolegend, 134717; 1:100) antibodies, followed by sheep anti-rat Dynabeads. Following magnetic separation, 1 × 10^5^ cells were seeded into collagen type I-coated glass-bottom dishes with cloning rings and cultured in DMEM/F-12 with GlutaMAX (Gibco, 10565018) supplemented with 10% FBS and 1% penicillin-streptomycin for 72 hours. The medium was then replaced with fresh medium containing cytokines in the following combinations: CXCL10 (1 μg/mL; PeproTech, 250-16), or TGF-β1 (1 ng/mL) alone; CXCL10 + TGF-β1 in combination. The cells were incubated with these cytokine treatments for 48 hours. Following cytokine stimulation, the cells were fixed with 4% PFA in PBS and processed for immunostaining.

### In vitro co-culture experiments of primary bone marrow-derived macrophages and primary lung fibroblasts

Bone marrow was harvested from femurs and tibiae of mice and filtered through a 100-μm cell strainer. The cells were washed with PBS and resuspended in red blood cell lysis buffer (Lonza, 10-548E) for 1 minute at room temperature. Following centrifugation, the cells were resuspended in DMEM/F-12 with GlutaMAX supplemented with 10% FBS, 1% penicillin-streptomycin, and M-CSF (100 ng/mL; kindly provided by Dr. Shinya Suzu, Kumamoto University). For establishment of bone marrow-derived macrophages, 5 × 10^5^ bone marrow cells were seeded into 35-mm petri dishes and cultured for 5 days. For co-culture experiments, 6 × 10^3^ primary lung fibroblasts isolated as described above were mixed with 6 × 10^4^ bone marrow-derived macrophages (1:10 ratio) and seeded into collagen type I-coated glass-bottom dishes with cloning rings in the presence or absence of CXCL10 (1 μg/mL). Cells were incubated for 48 hours, fixed with 4% PFA in PBS and processed for immunostaining.

### Immunofluorescence of culture cells

Fixed cells were blocked with 1% BSA in PBS with 0.1% Tween 20 for 1 hour at room temperature. Primary and secondary antibodies are listed in Table S1. Primary antibodies were incubated in the blocking buffer overnight at 4°C. Fluorescence-conjugated secondary antibodies were used at a 1:250 dilution in the blocking buffer (with Hoechst 33342 (1:500) or TO-PRO-3 (Thermo Fisher Scientific, T3605; 1:1000) when needed) and incubated overnight at 4°C. After washing in PBS-T, cells were mounted with Fluoromount-G.

### RNA isolation, reverse transcription (RT), and real time PCR analysis

Total RNA was isolated from the cells using the RNeasy Plus Micro Kit. Portions (50 ng) of the RNA were subjected to reverse transcription using ReverTra Ace qPCR RT Master Mix with gDNA Remover (Toyobo, FSQ-301). The resulting cDNA was subjected to real-time PCR analysis using Thunderbird Next SYBR qPCR Mix (Toyobo, QPX-201) and a LightCycler 96 instrument (Roche). The PCR conditions consisted of 40 cycles at 95°C for 5 s and 60°C for 10 s. The amount of each target mRNA was normalized to glyceraldehyde-3-phosphate dehydrogenase (*Gapdh*) mRNA. The sequences of the RT-qPCR primers were as follows: *Tubb3* forward: 5′-CAT GGA CAG TGT TCG GTC TG-3′ and *Tubb3* reverse: 5′-TTC CGC ACG ACA TCT AGG AC-3′. *Cxcl10* forward: CCG TCA TTT TCT GCC TCA TC-3′ and *Cxcl10* reverse: 5′-CTC GCA GGG ATG ATT TCA AG-3′. *Gapdh* forward: 5′-CAT CAC TGC CAC CCA GAA GAC TC-3′ and *Gapdh* reverse: 5′-ATG CCA GTG AGC TCC CGT TCA G-3′.

### scRNA-seq analysis of publicly available datasets from bleomycin-induced pulmonary fibrosis mouse models

Two publicly available scRNA-seq datasets of bleomycin-induced pulmonary fibrosis mouse model were analyzed; one dataset is from day 14 following bleomycin treatment in *Col1-GFP* reporter mice (Tsukui et al., 2020), while the other comprises multiple time points after bleomycin treatment in *Scube2-Cre^ERT2^; R26R-tdTomato* mice (Tsukui et al., 2024). The analysis was performed using the R package Seurat (Hao et al., 2021).

For the analysis of the *Col1-GFP* dataset, a cell-annotated Seurat object was provided by the Sheppard lab (UCSF). Using the cell annotation metadata, the pericyte population was extracted from the whole cell dataset and re-clustered following the Seurat standard workflow, including principal component analysis (PCA), followed by Uniform Manifold Approximation and Projection (UMAP) dimensional reduction (dims = 1:15) with resolution 0.8. Differentially expressed genes (DEGs) were identified using the FindMarkers function in Seurat. Average gene expressions were calculated using the AverageExpression function in Seurat. Gene Ontology (GO) enrichment analysis was performed using GOATOOLS Python library (Klopfenstein et al., 2018). Pathway activity inference was performed using decoupleR package (Badia-i-Mompel et al., 2022). UMAP plots were generated using either Seurat or scCustomize R package (Marsh et al., 2024). Kernel density estimate was calculated by Nebulosa (Alquicira-Hernandez and Powell, 2021). Volcano plots and heatmaps were generated using EnhancedVolcano and tidyheatmaps R packages, respectively (Blighe et al., 2018; Engler, 2024).

For the analysis of the *Scube2-Cre^ERT2^; R26R-tdTomato* dataset, the Seurat object was retrieved from the Gene Expression Omnibus (GEO) database under accession number GSE210341. For inflammatory fibroblasts and fibrotic fibroblasts, deposited cell annotation metadata was used and the number of cells in each fibroblast population was counted at each time point. For Tuj1^+^ pericytes, the total pericyte population was annotated using pericyte marker genes (*Higd1b*, *Cox4i2*, and *Ndufa4l2*), and Tuj1^+^ pericytes were subset based on *Tubb3* expression (> 0.5) and counted at each time point. Box plots were generated using Matplotlib Python library.

### scRNA-seq library preparation and sequencing for *Tubb3*^-/-^ mice

*Tubb3*^-/-^ mice and their wild-type littermates were treated with bleomycin, and lung tissues were harvested 14 days post-treatment. Two biological replicates were prepared for each genotype. The isolated lungs were dissociated and processed to obtain both whole cell samples and mural cell-enriched mesenchymal cell samples. For the whole cell samples, erythrocytes were removed by magnetic negative selection using a rat anti-mouse TER-119 (1:100) antibody followed by sheep anti-rat Dynabeads. For the mesenchymal cell samples, cells were enriched by magnetic negative selection using rat anti-mouse antibodies against CD31 (1:100), CD45 (1:100), EpCAM (1:100), and TER-19 (1:100), followed by sheep anti-rat Dynabeads. Following negative selection, both cell samples were counted and labeled with oligonucleotide tags for multiplexing using the 10x Genomics 3′ CellPlex Kit Set A (1000261). The labeled samples from all four mice were pooled according to sample type (whole cell or mesenchymal cell) and stained with DAPI (0.1 μg/mL) prior to cell sorting. For the pooled whole cell sample, live cells were sorted. For the pooled mesenchymal cell sample, MCAM^+^ PDGFRα^-^ (mural cells) and MCAM^-^ (non-mural cells) populations were sorted and mixed at a 1:1 ratio to achieve mural cell enrichment. Single-cell isolation, RNA capture, cDNA preparation, and library preparation were performed according to the manufacturer’s protocol (Chromium Next GEM Single Cell 3’ Reagent Kit, v3.1 chemistry, 10x Genomics). The libraries were sequenced on an Illumina NovaSeq 6000 S4 flow cell.

### scRNA-seq analysis of *Tubb3*^-/-^ mice

Sequencing reads were aligned to the mouse reference genome (mm10) using Cell Ranger (v7.1.0). Sample demultiplexing was performed using the "cellranger multi" pipeline. Further analysis was conducted using the Seurat R package. For quality control, cells with <200 or >7,500 genes, and cells with >15% mitochondrial genes were removed. Genes detected in fewer than 3 cells were also filtered out. Variable genes were identified using FindVariableFeatures function and integrated using RunFastMNN (Haghverdi et al., 2018) from the SeuratWrappers package. UMAP was used for dimensionality reduction. Cell clusters were identified using Seurat’s FindNeighbors and FindClusters functions (resolution = 0.4) from a total of 25,964 cells. Pericytes and vascular smooth muscle cells (VSMCs) were subset based on their specific markers and re-clustered following the standard Seurat workflow. UMAP plots were generated using either Seurat or scCustomize R packages. Average gene expressions were calculated using the AverageExpression function in Seurat.

For cell compositional analysis using scCODA (Büttner et al., 2021), an AnnData object was created using Scanpy (Wolf et al., 2018) Python library. Data was preprocessed with the same quality control parameters as above. Doublets were removed using scDblFinder (Germain et al., 2022). Following normalization and log transformation, principal component analysis was performed on highly variable genes. A neighborhood graph was constructed, followed by UMAP dimensionality reduction and Leiden clustering (resolution = 1.5). Major cell populations (fibroblasts, immune cells, endothelial cells, and epithelial cells) were subset and independently re-clustered using Leiden clustering with resolution parameters of 0.7, 2.0, 0.4, and 0.4, respectively. scCODA analysis was performed with a false discovery rate (FDR) threshold of 0.2, and results were visualized using scCODA’s built-in plotting functions.

### Statistical analysis

All data were collected from at least three independent experiments. The number of independent biological replicates is specified in the relevant figure legends where applicable. Statistical analyses were performed using GraphPad Prism (version 10.2.0). Data are presented as means ± SEM. Comparisons between two groups were performed using unpaired two-tailed Student’s *t*-tests, and comparisons among three groups were analyzed by one-way ANOVA followed by Tukey’s multiple comparisons test. A *P* value < 0.05 was considered statistically significant.

## Supplemental material

Supplemental Figure 1 – 5

Supplemental Table 1, 2

## Data availability

The original data presented in this study are included in the article. The scRNA-seq data generated in this study have been deposited in the Gene Expression Omnibus (GEO) under accession number GSE282025. Further inquiries can be directed to the corresponding author, Yoh-suke Mukouyama.

## Author contributions

RS designed and performed experiments, conducted data analysis, and wrote the manuscript. KI, TY, KO, and TS performed experiments. TT assisted with experiments. YT, KF, TI, CAC, MM, JBK, MS, TS, and DS provided experimental materials and procedures. Y-SM directed this study and wrote the manuscript.

## Acknowledgments

We thank K. Sakimura (Niigata University) for *Tubb3-Cre* mice; T. Arnold (UCSF) for *αSMA-Cre^ERT2^; ROSA-LSL-RFP* mice; S. Suzu (Kumamoto University) for M-CSF; P. Dagur and the staff of the NHLBI Flow Cytometry Core for operating the BD FACSAria Fusion cell sorter; Y. Li and the staff of the NHLBI DNA Sequencing and Genomics Core for operating the Illumina NovaSeq 6000 sequencer; D. Malide (NHLBI Light Microscopy Core) for image acquisition; T. Clark and the staff of NIH Bldg50 animal facility for assistance with mouse breeding and care; K. Gill for laboratory management, technical support and editorial advice on the manuscript; V. Sam, J Dawes, and S. Thacker for administrative assistance. Thanks also to members of the Laboratory of Stem Cell and Neurovascular Research for technical help and valuable discussion. Illustrations were created using BioRender. This work was supported by the Intramural Research Program of the National Heart, Lung, and Blood Institute, National Institutes of Health (HL005702-19 to Y.M.). R. Sato was supported by the Japan Society for the Promotion of Science (JSPS) NIH-KAITOKU. The authors have no conflicting financial interests.

