## Supplemental figures for "β-III tubulin identifies anti-fibrotic state of pericytes in pulmonary fibrosis"

**Figure S1.**

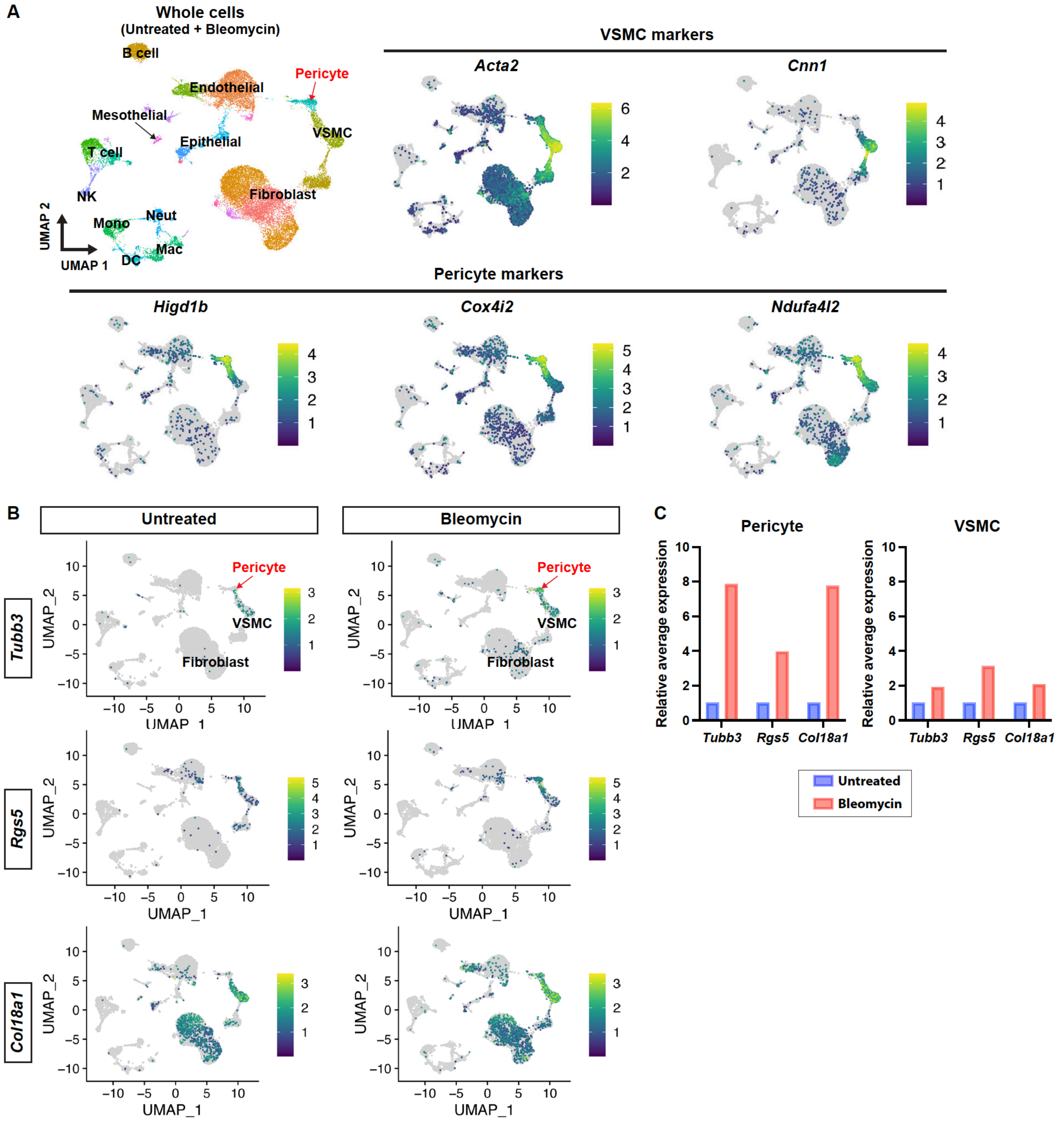


Figure S1. **Bleomycin-induced lung injury selectively upregulates *Tubb3* expression in lung pericytes.**

**(A)** UMAP plot of all cells from both untreated and bleomycin-treated samples, with cell type annotations, based on the scRNA-seq dataset from *Tsukui et al.*[21]. Expression of VSMC marker genes (*Acta2* and *Cnn1*) and pericyte marker genes (*Higd1b*, *Cox4i2*, and *Ndufa4l2*) is shown in UMAP plots. **(B)** UMAP plots split by treatment conditions (untreated and bleomycin-treated) showing expression of activated pericyte marker genes (*Tubb3*, *Rgs5*, and *Col18a1*) across all cells. **(C)** Comparison of the average expression of activated pericyte marker genes in pericyte (left) and VSMC (right) populations between untreated (blue) and bleomycin-treated (red) samples. Y-axis shows fold change in average expression relative to untreated samples (set to 1). VSMC: vascular smooth muscle cell; NK: natural killer cell; Mono: monocyte; Neut: neutrophil; Mac: macrophage; DC: dendritic cell.

Figure S2.

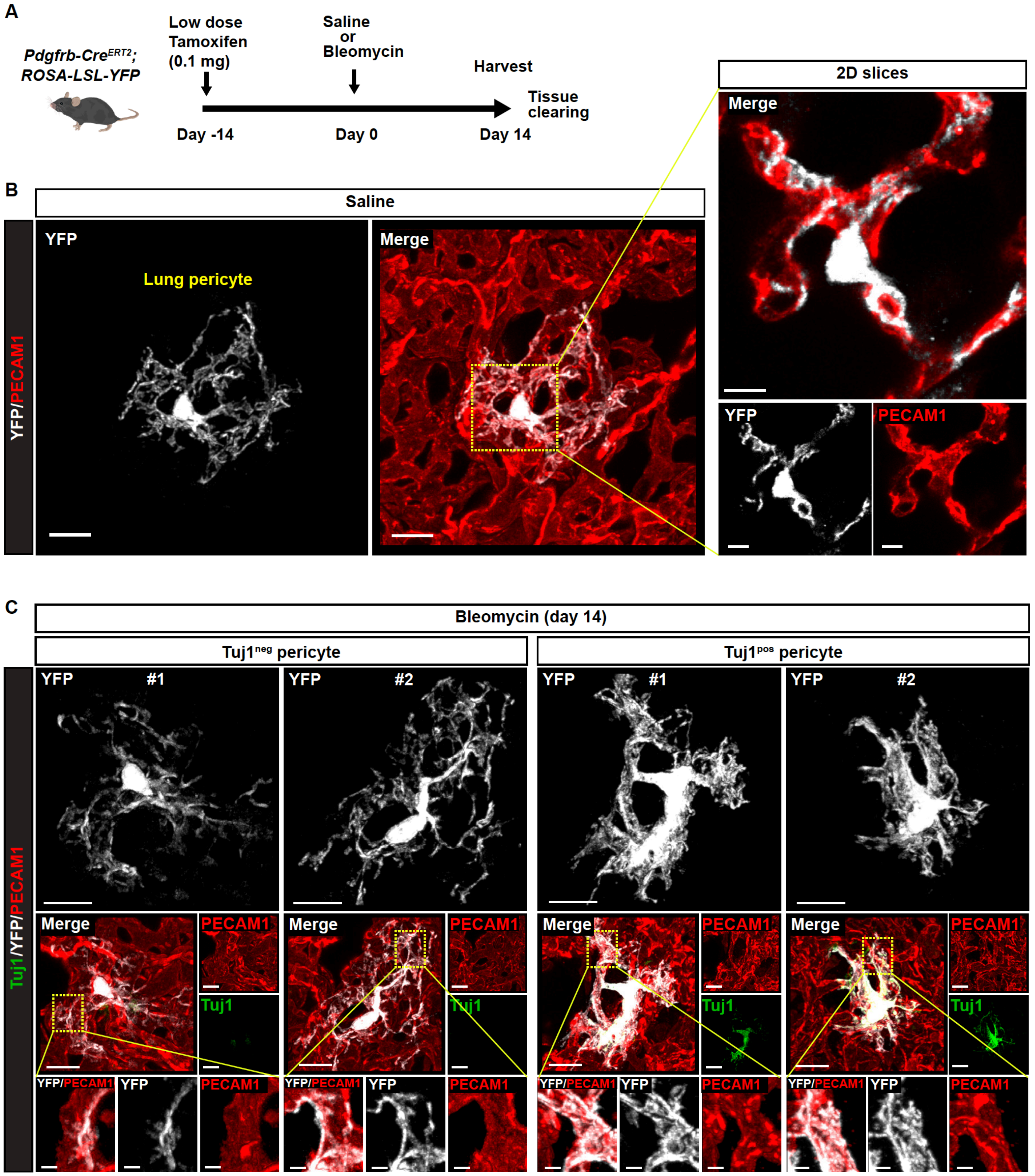


Figure S2. **Cellular morphology of single lung pericytes in homeostatic and bleomycin-induced fibrotic conditions.**

**(A)** Schematic illustration of mosaic pericyte labeling using *Pdgfrb-Cre$^{ERT2}$; ROSA-LSL-YFP* mice. A single low-dose tamoxifen (0.1 mg) was administered before saline or bleomycin treatment, with lung tissues harvested on day 14. **(B)** Immunostaining of tissue-cleared lungs from saline-treated *Pdgfrb-Cre$^{ERT2}$; ROSA-LSL-YFP* mice with YFP (gray) and PECAM1 (red). Attenuated-maximum intensity projection (attenuated-MIP) was used to visualize 3D lung vascular structures. Right panels: magnified 2D slice images of boxed regions in the left panel. Scale bars, 10 µm (left panels); 5 µm (right panels). **(C)** Immunostaining of tissue-cleared lungs from bleomycin-treated *Pdgfrb-Cre$^{ERT2}$; ROSA-LSL-YFP* mice with YFP (gray), Tuj1 (green), and PECAM1 (red). Representative images of Tuj1$^{neg}$ and Tuj1$^{pos}$ pericytes are shown (two cells per phenotype). Boxed regions in attenuated-MIP rendered images of Tuj1$^{neg}$ and Tuj1$^{pos}$ pericytes (middle panels) are magnified as corresponding 3D cropped images on the bottom panels. Scale bars, 10 µm; 2 µm (3D cropped).

**Figure S3.**

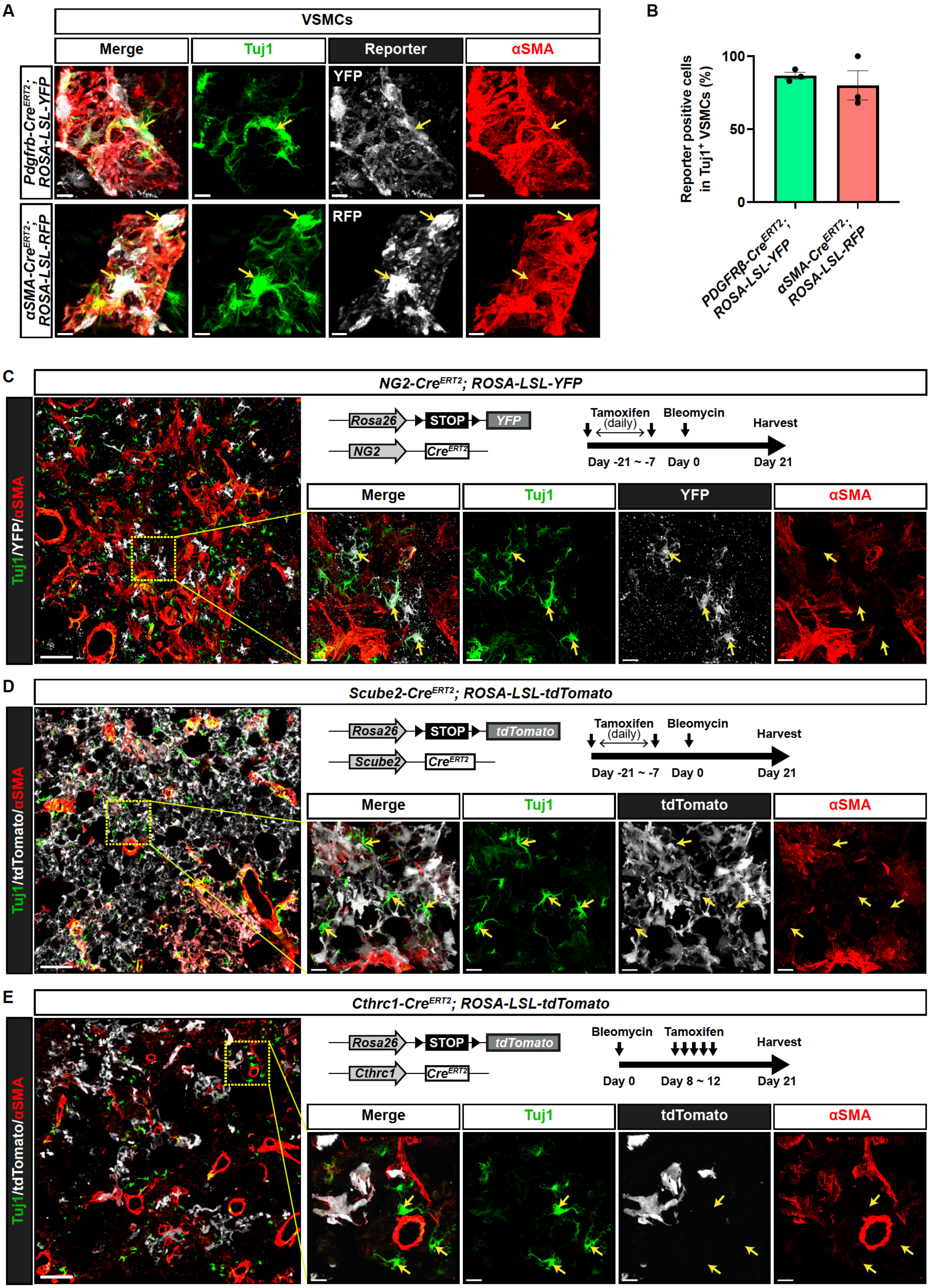
A
VSMCs
Merge
Tuj1
Reporter
αSMA
Pdgfrb-CreERT2; ROSA-LSL-YFP
YFP
αSMA-CreERT2; ROSA-LSL-RFP
RFP
B
Reporter positive cells in Tuj1+ VSMCs (%)
100
50
0
PDGFRβ-CreERT2; ROSA-LSL-YFP
αSMA-CreERT2; ROSA-LSL-RFP
C
NG2-CreERT2; ROSA-LSL-YFP
Tuj1/YFP/αSMA
Rosa26
STOP
YFP
NG2
CreERT2
Tamoxifen (daily)
Bleomycin
Harvest
Day -21 ~ -7
Day 0
Day 21
Merge
Tuj1
YFP
αSMA
D
Scube2-CreERT2; ROSA-LSL-tdTomato
Tuj1/tdTomato/αSMA
Rosa26
STOP
tdTomato
Scube2
CreERT2
Tamoxifen (daily)
Bleomycin
Harvest
Day -21 ~ -7
Day 0
Day 21
Merge
Tuj1
tdTomato
αSMA
E
Cthrc1-CreERT2; ROSA-LSL-tdTomato
Tuj1/tdTomato/αSMA
Rosa26
STOP
tdTomato
Cthrc1
CreERT2
Bleomycin
Tamoxifen
Harvest
Day 0
Day 8 ~ 12
Day 21
Merge
Tuj1
tdTomato
αSMA

Figure S3. **Lineage-tracing experiments revealed that neither VSMCs nor fibroblasts contribute to Tuj1$^+$ pericytes.**

**(A and B)** Lineage-tracing experiments of VSMCs using *Pdgfrb-Cre$^{ERT2}$; ROSA-LSL-YFP* and *αSMA-Cre$^{ERT2}$; ROSA-LSL-RFP* mice. **(A)** Immunostaining of bleomycin-treated lungs from *Pdgfrb-Cre$^{ERT2}$; ROSA-LSL-YFP* (upper panels) and *αSMA-Cre$^{ERT2}$; ROSA-LSL-RFP* (lower panels) mice, with Tuj1 (green), lineage-specific reporter protein (gray), and αSMA (red). Scale bars, 5 μm. **(B)** Proportion of YFP- or RFP-expressing cells within Tuj1$^+$ VSMCs. **(C)** Lineage-tracing experiments using *NG2-Cre$^{ERT2}$; ROSA-LSL-YFP* mice. The Cre-mediated excision was induced by administrating tamoxifen for 14 days prior to initiating bleomycin treatment, with lung tissues harvested on day 21. Immunostaining of bleomycin-treated lungs with Tuj1 (green), YFP (gray), and αSMA (red). Boxed region in the left panel is magnified in the right panels. Arrows indicate Tuj1$^+$ pericytes. Scale bars, 100 μm (left); 15 μm (magnified views). **(D)** Lineage-tracing experiments using *Scube2-Cre$^{ERT2}$; ROSA-LSL-tdTomato* mice. The Cre-mediated excision was induced by administrating tamoxifen for 14 days prior to initiating bleomycin treatment, with lung tissues harvested on day 21. Immunostaining of bleomycin-treated lungs with Tuj1 (green), tdTomato (gray), and αSMA (red). Boxed region in the left panel is magnified in the right panels. Arrows indicate Tuj1$^+$ pericytes. Scale bars, 100 μm (left); 15 μm (magnified views). **(E)** Lineage-tracing experiments using *Cthrc1-Cre$^{ERT2}$; ROSA-LSL-tdTomato* mice. The Cre-mediated excision was induced by administrating tamoxifen for 5 days, starting 8 days post-bleomycin treatment, with lung tissues harvested on day 21. Immunostaining of bleomycin-treated lungs with Tuj1 (green), tdTomato (gray), and αSMA (red). Boxed region in the left panel is magnified in the right panels. Arrows indicate Tuj1$^+$ pericytes. Scale bars, 100 μm (left); 15 μm (magnified views).

**Figure S4.**

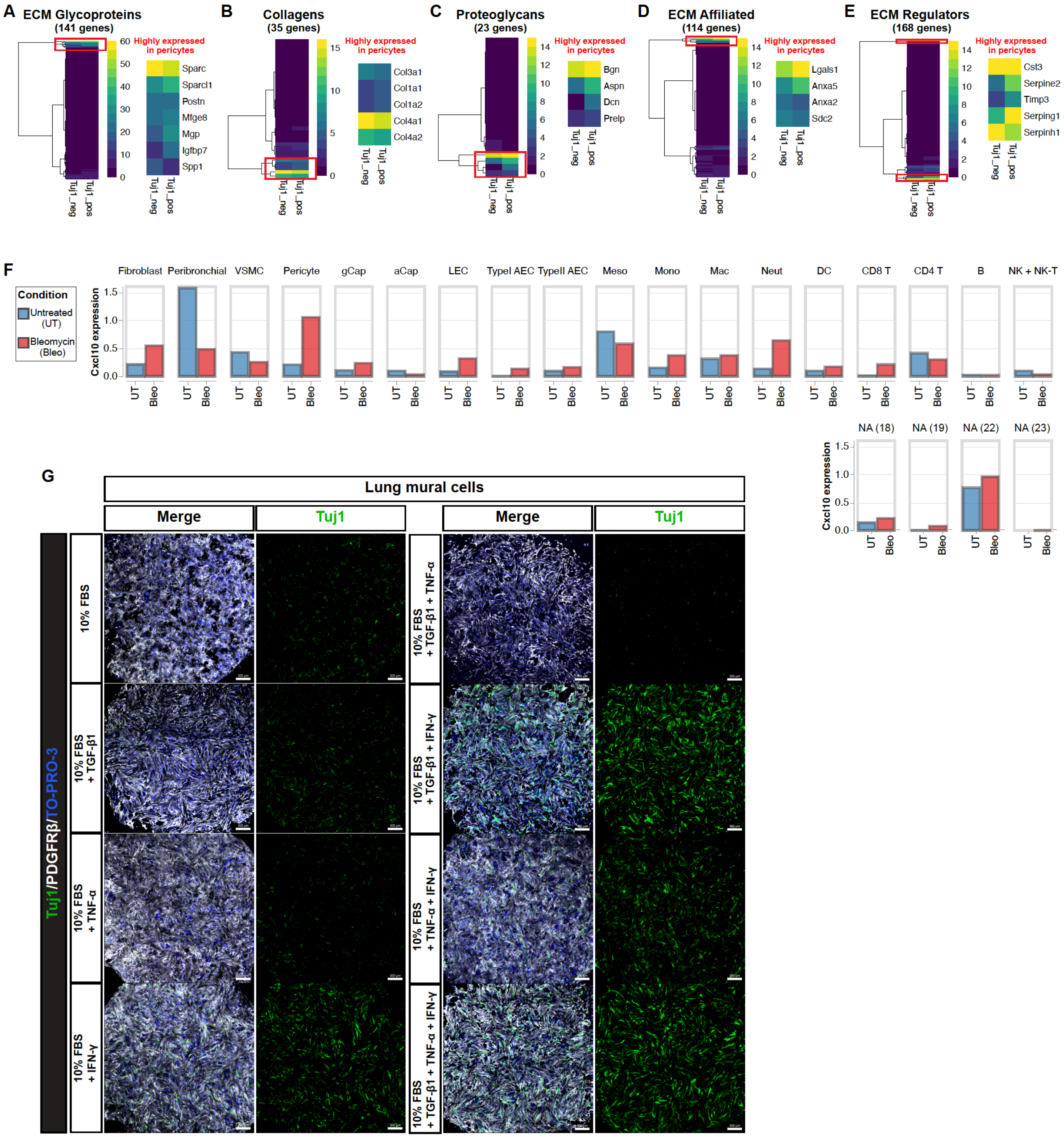


Figure S4. **Gene expression profiles and upstream signals regulating Tuj1 in lung pericytes.** **(A–E)** Heatmap of the average expression levels for matrisome genes in Tuj1$^{neg}$ and Tuj1$^{pos}$ pericytes from bleomycin-treated lung samples, based on scRNA-seq data[21]. Matrisome categories displayed: (A) ECM Glycoproteins, (B) Collagens, (C) Proteoglycans, (D) ECM Affiliated, and (E) ECM Regulators. For each category, a highly expressed gene cluster (highlighted by a red rectangle) is enlarged on the right, with corresponding gene names shown. ECM: extracellular matrix. **(F)** Average expression of *Cxcl10* in various cell types from untreated (blue) and bleomycin-treated (red) samples of the scRNA-seq dataset from *Tsukui et al.*[21]. Cell type annotations are based on the original publication. UT: untreated; Bleo: bleomycin-treated; Peribronchial: peribronchial fibroblast; VSMC: vascular smooth muscle cell; gCap: general capillary cell; aCap: aerocyte; LEC: lymphatic endothelial cell; TypeI AEC: type I alveolar epithelial cell; TypeII AEC: type 2 alveolar epithelial cell; Meso: mesothelial cell; Mono: monocyte; Mac: macrophage; Neut: neutrophil; DC: dendritic cell; CD8 T: CD8 T cell; CD4 T: CD4 T cell; B: B cell; NK + NK-T: natural killer cell + natural killer T cell; NA: not applicable. **(G)** Immunostaining of cultured lung mural cells with PDGFRβ (gray), Tuj1 (green), and TO-PRO-3 (blue). The freshly isolated lung mural cells were cultured for 48 hours in growth medium containing 10% fetal bovine serum (FBS), either alone or supplemented with IFN-γ, TGF-β1 or TNF-α, pairwise combinations of these cytokines, or all these cytokines together. Note that the combination of IFN-γ and TGF-β1 effectively induced Tuj1 expression in lung mural cells in the primary culture. Scale bars, 300 μm.

**Figure S5.**

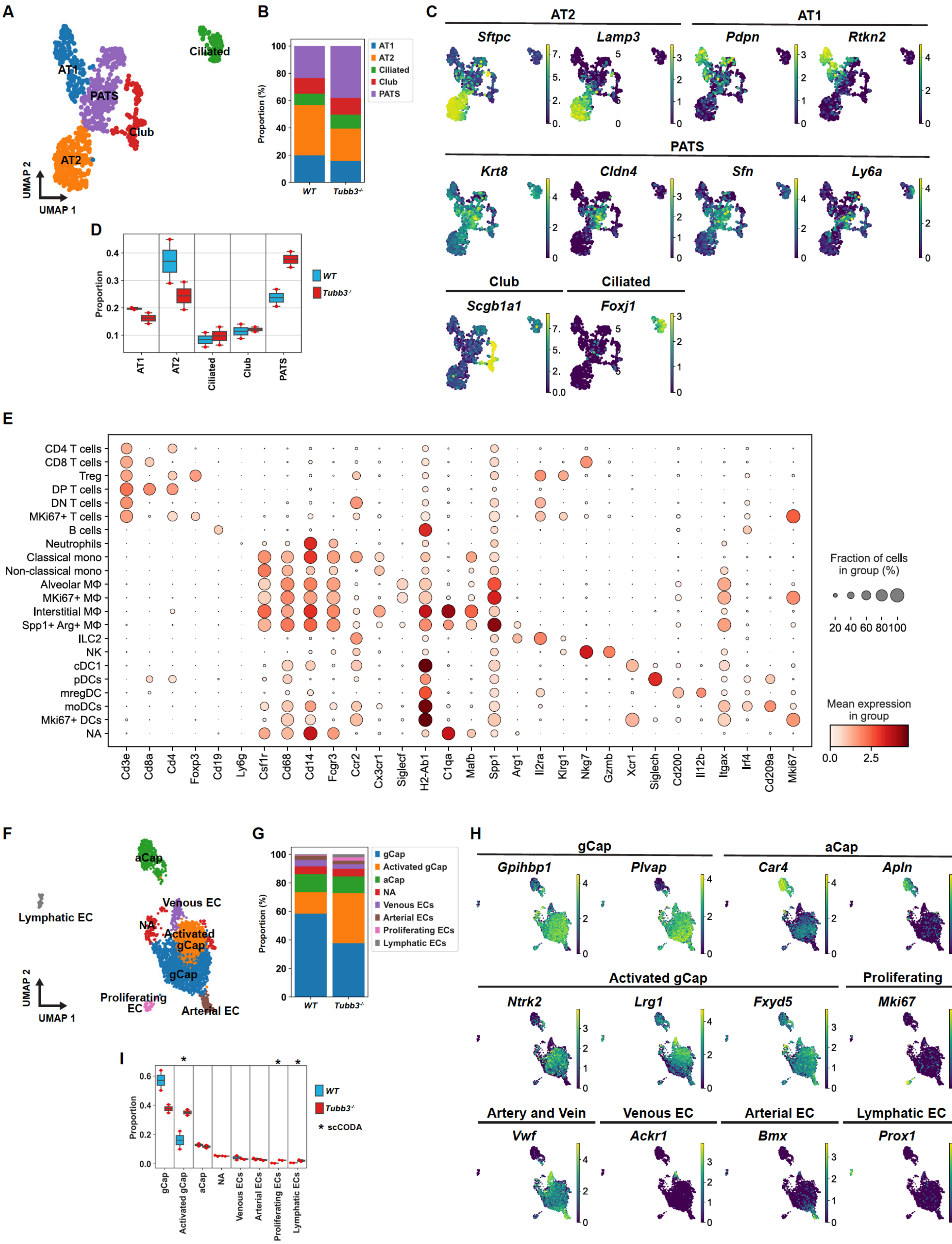

A
Ciliated
AT1
PATS
Club
AT2
UMAP 2
UMAP 1
B
AT1
AT2
Ciliated
Club
PATS
Proportion (%)
WT
Tubb3-/-
C
AT2
AT1
Sftpc
Lamp3
Pdpn
Rtkn2
PATS
Krt8
Cldn4
Sfn
Ly6a
Club
Ciliated
Scgb1a1
Foxj1
D
Proportion
WT
Tubb3-/-
AT1
AT2
Ciliated
Club
PATS
E
CD4 T cells
CD8 T cells
Treg
DP T cells
DN T cells
MKi67+ T cells
B cells
Neutrophils
Classical mono
Non-classical mono
Alveolar MΦ
MKi67+ MΦ
Interstitial MΦ
Spp1+ Arg+ MΦ
ILC2
NK
cDC1
pDCs
mregDC
moDCs
Mki67+ DCs
NA
Cd3e
Cd8a
Cd4
Foxp3
Cd19
Ly6g
Csf1r
Cd68
Cd14
Fcgr3
Ccr2
Cx3cr1
Siglecf
H2-Ab1
C1qa
Mafb
Spp1
Arg1
Il2ra
Klrg1
Nkg7
Gzmb
Xcr1
Siglech
Cd200
Il12b
Itgax
Irf4
Cd209a
Mki67
Fraction of cells in group (%)
20 40 60 80 100
Mean expression in group
0.0 2.5
F
aCap
Lymphatic EC
Venous EC
NA
Activated gCap
gCap
Proliferating EC
Arterial EC
UMAP 2
UMAP 1
G
gCap
Activated gCap
aCap
NA
Venous ECs
Arterial ECs
Proliferating ECs
Lymphatic ECs
Proportion (%)
WT
Tubb3-/-
H
gCap
aCap
Gpihbp1
Plvap
Car4
Apln
Activated gCap
Proliferating
Ntrk2
Lrg1
Fxyd5
Mki67
Artery and Vein
Venous EC
Arterial EC
Lymphatic EC
Vwf
Ackr1
Bmx
Prox1
I
Proportion
WT
Tubb3-/-
* scCODA
gCap
Activated gCap
aCap
NA
Venous ECs
Arterial ECs
Proliferating ECs
Lymphatic ECs

Figure S5. **Cell compositional analysis of epithelial cells and endothelial cells.**

**(A–D)** Cell compositional analysis of epithelial populations from scRNA-seq data of *WT* and *Tubb3*$^{-/-}$ mice 14 days after bleomycin treatment. UMAP plot of epithelial cells showing 5 subtypes (A) as defined by marker genes (C). **(B and D)** Proportions of each subtype relative to the total epithelial cells in *WT* and *Tubb3*$^{-/-}$ mice are shown as stacked barplot (B) and boxplot (D). No significant changes were observed by scCODA analysis in panel (D). AT1: alveolar type I cell; AT2: alveolar type II cell; PATS: pre-alveolar type-1 transitional cell state cell; Club: club cell; Ciliated: ciliated cell. **(E)** Dot plot showing expression of marker genes for immune cell subtypes from scRNA-seq data of bleomycin-treated lungs (day 14) from *WT* and *Tubb3*$^{-/-}$ mice. **(F–I)** Cell compositional analysis of endothelial populations from scRNA-seq data of *WT* and *Tubb3*$^{-/-}$ mice 14 days after bleomycin treatment. UMAP plot of endothelial cells showing 7 subtypes (F) as defined by marker genes (H). **(G and I)** Proportions of each subtype relative to the total endothelial cells in *WT* and *Tubb3*$^{-/-}$ mice are shown as stacked barplot (G) and boxplot (I). Statistically credible changes, as tested by scCODA, are noted with an * in panel (I). gCap: general capillary cell; aCap: aerocyte; EC: endothelial cell; NA: not applicable.

**Table S1. Lists of primary and secondary antibodies for immunostaining**

| Primary antibody | Company | Catalog number | Host | Dilution |
|---|---|---|---|---|
| Tuj1-Alexa 488 | Biolegend | 801203 | Mouse | 1:200 |
| Tuj1-Alexa 647 | Biolegend | 801209 | Mouse | 1:200 |
| aSMA-FITC | Millipore Sigma | F3777 | Mouse | 1:500 |
| aSMA-Cy3 | Millipore Sigma | C6198 | Mouse | 1:500 |
| PECAM1 | Chemicon | MAB1398z | hamster | 1:300 |
| PDGFRb | eBioscience | 14-1402-82 | Rat | 1:100 |
| PDGFRβ | R&D | AF1042-SP | Goat | 1:100 |
| NG2 | Millipore Sigma | AB5320 | Rabbit | 1:200 |
| MCAM-Alexa 647 | Biolegend | 134717 | Rat | 1:100 |
| GFP | Thermo Fisher | A-11122 | Rabbit | 1:1000 |
| GFP | Abcam | ab6673 | Goat | 1:500 |
| DsRed | TaKaRa | 632496 | Rabbit | 1:1000 |
| Collagen 1 | Millipore Sigma | AB765P | Rabbit | 1:200 |
| Podoplanin-Alexa 594 | Biolegend | 127414 | Syrian hamster | 1:100 |
| Podoplanin-APC | Biolegend | 127410 | Syrian hamster | 1:100 |
| TrkB | R&D | AF1494-SP | Goat | 1:200 |
| PDGFRa | R&D | AF1062-SP | Goat | 1:200 |
| CD45 | eBioscience | 14-0451-85 | Rat | 1:500 |
| CD68-Alexa 488 | Biolegend | 137011 | Rat | 1:100 |
| CD68-Alexa 594 | Biolegend | 137020 | Rat | 1:100 |
| Arginase-PE | Thermo Fisher | 12-3697-80 | Rat | 1:200 |
| SPP1 | R&D | AF808-SP | Goat | 1:400 |
| Ki67-Alexa 488 | BD | 558616 | Mouse | 1:100 |
| PGP9.5 | Abcam | ab15503 | Rabbit | 1:200 |
| TH | Millipore Sigma | AB152 | Rabbit | 1:200 |

| | | | | |
|---|---|---|---|---|
| CSF1R | Santa Cruz | sc-692 | Rabbit | 1:400 |

| Secondary antibody | Company | Catalog number | Host | Dilution |
|---|---|---|---|---|
| anti Armenian hamster-Alexa 488 | Jackson ImmunoResearch | 127-545-160 | Goat | 1:250 |
| anti Armenian hamster-Cy3 | Jackson ImmunoResearch | 127-165-160 | Goat | 1:250 |
| anti Armenian hamster-Alexa 647 | Jackson ImmunoResearch | 127-605-160 | Goat | 1:250 |
| anti Goat-Alexa 488 | Jackson ImmunoResearch | 705-547-003 | Donkey | 1:250 |
| anti Goat-Cy3 | Jackson ImmunoResearch | 705-167-003 | Donkey | 1:250 |
| anti Goat-Alexa 594 | Jackson ImmunoResearch | 705-585-147 | Donkey | 1:250 |
| anti Rabbit-Alexa 488 | Invitrogen | A11034 | Goat | 1:250 |
| anti Rabbit-Cy3 | Jackson ImmunoResearch | 711-165-152 | Donkey | 1:250 |
| anti Rabbit-Alexa 647 | NanoTag | N2404 | Alpaca | 1:250 |
| anti Rat-Cy3 | Jackson ImmunoResearch | 712-166-153 | Donkey | 1:250 |

**Table S2. List of hybridization probes for *Cxcl10***

| Probe | Sequence |
|---|---|
| Cxcl10_A1P1O | AGAGAATCATACGTtaCTAGGGAGGACAAGGAGGGTGTGGG |
| Cxcl10_A1P1E | GGTAAACTTAGAACTGACGAGCCTGaaCGTCAACGACAAGC |
| Cxcl10_A1P2O | AGAGAATCATACGTtaGCGTCGCACCTCCACATAGCTTACA |
| Cxcl10_A1P2E | CAGCCTGGGCATGGCACATGGTGAAaaCGTCAACGACAAGC |
| Cxcl10_A1P3O | AGAGAATCATACGTtaTTGAGCGAGGACTCAGACCAGCCCT |
| Cxcl10_A1P3E | CAGAGCTAGGACAGCCATCCCAGCCaaCGTCAACGACAAGC |
| Cxcl10_A1P4O | AGAGAATCATACGTtaGTCTCAGGACCATGGCTTGACCATC |
| Cxcl10_A1P4E | AGAATTCTTGCTTCGGCAGTTACTTaaCGTCAACGACAAGC |
| Cxcl10_A1P5O | AGAGAATCATACGTtaCCACGGCTGGTCACCTTTCAGAAGA |
| Cxcl10_A1P5E | CTGCAGGAGGAGTAGCAGCTGATGTaaCGTCAACGACAAGC |
| Cxcl10_A1P6O | AGAGAATCATACGTtaGGTAAAGGGGAGTGATGGAGAGAGG |
| Cxcl10_A1P6E | AGGGCAATTAGGACTAGCCATCCACaaCGTCAACGACAAGC |
| Cxcl10_A1P7O | AGAGAATCATACGTtaGGTGTGTGCGTGGCTTCACTCCAGT |
| Cxcl10_A1P7E | TCTGCTGTCCATCCATCGCAGCACCaaCGTCAACGACAAGC |
| Cxcl10_A1P8O | AGAGAATCATACGTtaCACTGGCCCGTCATCGATATGGATG |
| Cxcl10_A1P8E | TTCAAGCTTCCCTATGGCCCTCATTaaCGTCAACGACAAGC |
| Cxcl10_A1P9O | AGAGAATCATACGTtaATCCCTTGAGTCCCACTCAGACCCA |
| Cxcl10_A1P9E | TTGCAGCGGACCGTCCTTGCGAGAGaaCGTCAACGACAAGC |
| Cxcl10_A1P10O | AGAGAATCATACGTtaTTGGGTTCATGGTGCTGATGGGGAG |
| Cxcl10_A1P10E | GGATGAGGCAGAAAATGACGGCAGCaaCGTCAACGACAAGC |
